# Convergent evolution through independent rearrangements in the primate amylase locus

**DOI:** 10.1101/2025.08.14.670395

**Authors:** Charikleia Karageorgiou, Petar Pajic, Stefan Ruhl, Omer Gokcumen

## Abstract

Structurally complex genomic regions can foster evolutionary convergence by repeatedly generating gene duplications that yield similar expression patterns and traits across lineages. Focusing on the primate amylase locus, we leveraged high-quality genome assemblies from 53 primate species and multi-tissue transcriptomes from Old World monkeys to reconstruct the evolutionary history of recurrent duplications. We show that lineage-specific long terminal repeat retrotransposon insertions may be associated with initial structural instability, while subsequent duplications are primarily driven by non-allelic homologous recombination. Independent duplications in rhesus macaques, olive baboons, and great apes produced distinct amylase copies with convergent expression in pancreas and salivary glands and signals of episodic diversifying selection, consistent with emerging functional divergence. Our analyses indicate that a ancestral gene with dual pancreas and salivary expression in Catarrhini duplicated in great apes, facilitating subfunctionalization and regulatory rewiring. These findings illuminate how modular structural and regulatory variation drives evolutionary innovation and molecular convergence.

## INTRODUCTION

Convergent evolution, the independent emergence of similar traits in distinct lineages, has long intrigued evolutionary biologists seeking to understand how similar phenotypes arise from different genetic starting points^1,2^ Recent advances in long-read sequencing have allowed us to characterize structurally complex genomic regions accurately at the nucleotide resolution, revealing that these regions frequently undergo recurrent structural variation^3–6^ including gene duplications, that can drive major shifts in biological function^7–11^ Integrating these insights, an emerging model posits that structurally complex loci, through repeated gene duplications and regulatory rewiring, can serve as substrates for molecular convergence ^5,12,13^ n particular, such loci may produce similar spatial expression patterns of gene families n phylogenetically distant lineages. Yet, the specific mutational mechanisms that give rise to these duplications, the evolutionary forces shaping nucleotide variation among paralogs, and the processes by which regulatory elements are reshuffled during structural rearrangements remain largely unresolved. These questions are at the heart of understanding how structural and regulatory complexity contributes to evolutionary innovation and the repeated emergence of similar traits across lineages.

The amylase locus is one of the most intriguing structurally complex regions in mammalian genomes, notable for its exceptionally rapid structural evolution. It is one of the fastest-evolving loci in the human genome, despite being essential in starch metabolism^14^ Amylase in mammals is primarily expressed in the pancreas. However, in some lineages, the gene has undergone a regulatory shift to include expression in the salivary glands^15–17^ Variation in amylase copy number has been proposed to relate to dietary starch intake in several lineages. In humans, salivary amylase (*AMY1*) copy number shows population-level associations with starch-rich diets^3,18,19^ albeit the evolutionary and functional interpretation of this relationship remains an area of active investigation^20^ Similar patterns of amylase gene duplications and concordant expression shifts of the duplicated copies have evolved multiple times, independently in different mammalian lineages, thus suggesting convergent mechanisms in response to dietary shifts^17,19,21,22^ However, the mutational mechanisms underlying the independent gene duplications in the amylase locus and proximate regulatory sequences remain unexplored. Investigating these processes, using this locus as a model, could provide key insights into fundamental aspects of genomic evolution, including neofunctionalization, subfunctionalization, and gene expression dosage regulation.

In humans, saliva amylase is the most abundantly secreted enzyme in the oral cavity^23^ with expression levels *6-8-*fold higher than in other great apes^18,24,25^ This heightened expression was considered a human-specific trait^15,17,18^ However, across the primate phylogeny, other species also exhibit high salivary amylase expression, including Old World monkeys^26^ and capuchins^17^ The prevailing model suggests that the ancestral amylase gene was expressed initially in the pancreas, and was duplicated independently in different primate lineages, where some duplications acquired expression in the salivary glands. In apes, where this process has been best studied, the shift to salivary expression of one *AMY1* duplicate has been attributed to the insertion of an endogenous retroviral element (ERV) upstream of *AMY1*^17,27–29^ In humans, additiona *AMY1* duplications led to increased saliva expression levels^30^ In other nonhuman primates, the genetic and regulatory mechanisms underlying salivary-gland-specific amylase expression remain unknown.

Therefore, we compared the evolutionary history of the amylase locus in great apes and Old World monkeys, offering an ideal framework for investigating how rapidly evolving, structurally complex regions can give rise to convergent gene expression patterns. Specifically, we asked whether the recurrent gain of salivary gland expression of *AMY* in these lineages is driven by shared mutational mechanisms and regulatory shifts, or by distinct molecular events that lead to similar outcomes. To address this, we leverage 244 high-quality primate genome assemblies and transcriptome data from pancreas, liver, and salivary glands of rhesus macaques (*Macaca mulatta*) and olive baboons (*Papio anubis*). Using this data, we identified features of the amylase locus associated with salivary gland expression, characterize the mutational processes underlying lineage-specific gene duplications, and examine how structural and regulatory variation is linked to *AMY* function and expression. In doing so, our study not only advances understanding of amylase locus evolution and regulation in primates but also provides a broader model for investigating how gene duplications in complex genomic regions can drive convergent evolution of tissue-specific expression.

## RESULTS

### Ancestral and recurrent independent duplications shape primate amylase genetic structural diversity

Previous studies have documented extensive structural variation within the human amylase locus, identifying multiple independent copy number changes and inversions^3,4^ To determine if similar structural complexity extends across non-human primates, we analyzed 244 primate genomes, successfully curating continuous contigs spanning the amylase locus in 53 species (total 69 genomes **Table S1 & 2;** see **Methods**). For 16 species, multiple high-quality genome assemblies (ranging from 2 to 4 genomes per species) exist, allowing us to assess within-species variation (**Table S3**). In the remaining 174 genomes, the amylase locus could not be completely resolved within a single contig, underscoring the challenges of assembling this complex region, even with high-quality long-read data. Nevertheless, the 69 curated genomes provided a robust framework for reconstructing *AMY* structural variation across primates, documenting the expansions and contractions of the copy number variation of *AMY* genes, and predicting the ancestral structural states (**Figure 1A, Table S4,** see **Methods**).

**Figure 1.**
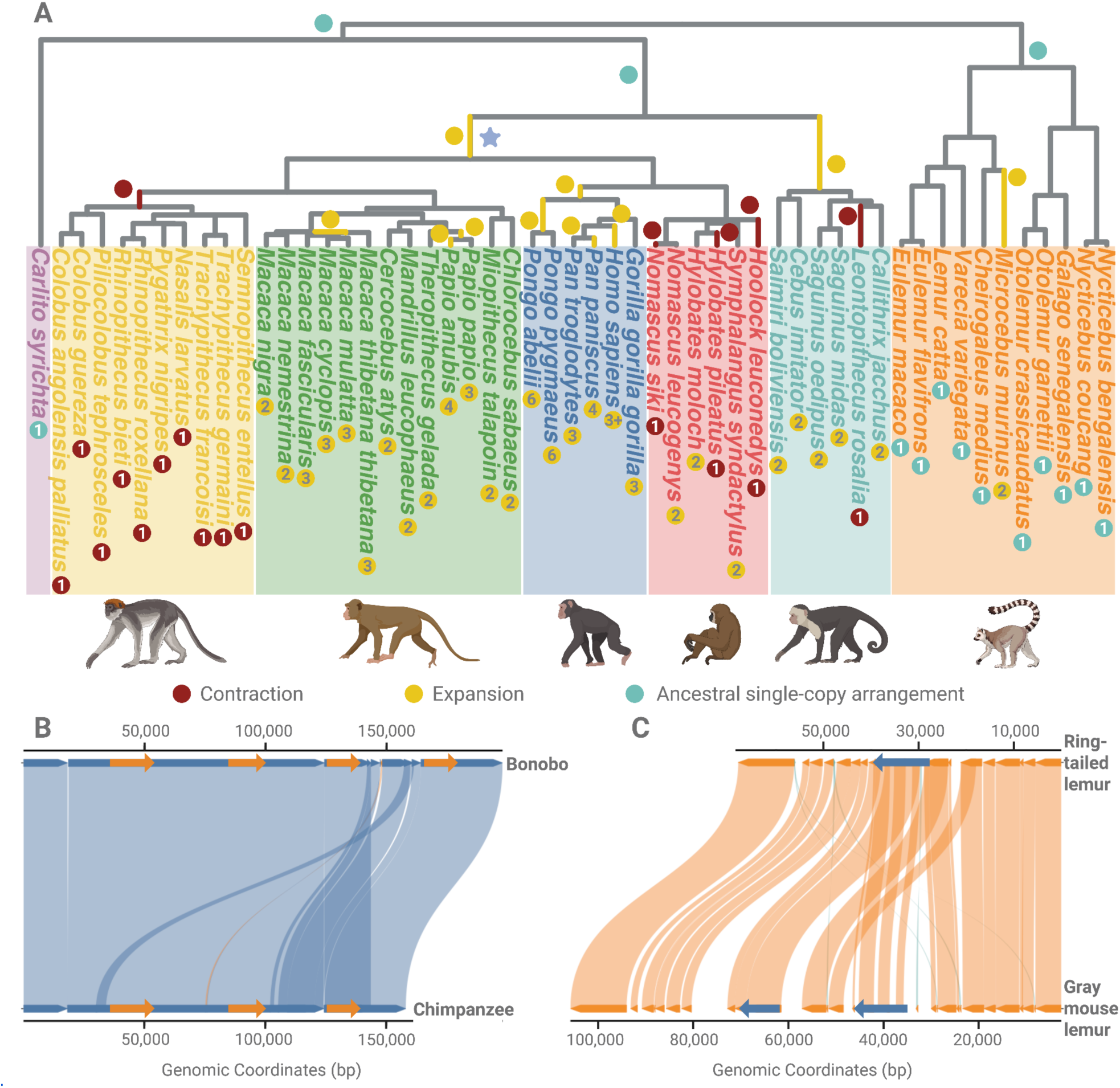
Structural evolutionary history of the amylase locus across primates. (A) Contractions and expansions in the amylase locus reconstructed from 69 high-quality genomes representing 53 primate species (**Table S2 & S4**). Lineages are color-coded by clade: purple for tarsiers (outgroup), yellow for leaf-eating monkeys, green for Old World monkeys, dark blue for great apes, red for lesser apes (gibbons), cyan for New World monkeys, and orange for lemurs. Red dots indicate independent contractions in the amylase locus. Yellow dots indicate expansions. Cyan dots mark lineages retaining the ancestral single-copy configuration, inferred as the ancestral primate state Independent contraction events are observed in the gibbon, leaf-eating monkey, and New World monkey genera. Numbers inside the dots give the total number of *AMY* gene copies detected in each species (per haploid genome). (B) Synteny comparison between bonobo (*Pan paniscus*) and chimpanzee (*Pan troglodytes*) at the amylase locus. The copy number increase in bonobo, previously reported^18^ is shown here to involve a chimeric duplication. The 5′ flanking region of the duplicated segment resembles the downstream region of *AMY1* The internal genic region corresponds to a full duplication of the *AMY1* coding sequence. The 3′ flanking region aligns with the upstream flanking region of *AMY2A* This chimeric architecture is consistent with a nonallelic homologous recombination mechanism. (C) Comparison of the amylase locus in ring-tailed lemur (*Lemur catta*) and gray mouse lemur (*Microcebus murinus*). The gray mouse lemur harbors a tandem duplication of the amylase gene including both upstream and downstream flanking regions. This is the only duplication identified among the lemur species analyzed.

Our results confirmed previous work^17,27^ that identified an ancestral duplication of the amylase locus in the Catarrhini lineage (Catarrhini: the parvorder comprising Old World monkeys and apes; ndicated by purple star in **Figure 1A**), lineage-specific duplications in New World monkeys, and a loss of a single *AMY* gene in leaf-eating monkeys^17^ The recently available high-quality genome assemblies analyzed in this study provide sequence-level resolution of lineage-specific copy number variations and ed to the discovery of novel duplication events. These include a burst of duplications in the orangutan lineage, a previously noted^18^ but uncharacterized copy number increase in bonobos relative to chimpanzees (**Figure 1B**), and recurrent independent duplications within Old World monkey genera, as described below. In addition, we found copy number variation among lesser ape species (i.e., in gibbons) (**Figure 1A; Figure S1; Table S5**), yet it remains unclear whether this variation reflects an ancestral loss followed by lineage-specific gains, independent lineage-specific losses, or incomplete lineage sorting (**Figure S2**), posing an intriguing question for future research Lemurs provide a ideal outgroup for studying the rest of the primate phylogeny. We found that all (n=11) but one of the analyzed lemur species harbor a single *AMY* gene per haploid genome, a structure shared with several non-lemur primates, suggesting that this configuration likely represents the ancestral amylase haplotype in primates. One exception is *Microcebus murinus* which carries an additional *AMY* copy (**Figure 1C**). Taken together, our findings illustrate a complex evolutionary history of the amylase locus characterized by recurrent independent duplications and losses, thus contributing a remarkable structural diversity in primates. These patterns are consistent with the amylase region behaving as a mutational hotspot across primates as defined by^31–33^ More genomes from additional species and more comprehensive dietary data would be required to robustly test the relationship between diet and *AMY* copy number in primates. Our study sets the stage to more comprehensively analyze the evolution of functional variation within a hotspot of genomic variation within closely related primate species with distinct dietary habits.

### Reconstruction of the amylase duplications in catarrhini

To further elucidate the mutational basis of *AMY* structural variation in primates, we closely examined the rhesus macaque and olive baboon genomes, contrasting their evolutionary histories with recent findings from human haplotypes^4^ Previous studies leveraged Old World monkeys as an outgroup to explore *AMY* copy number expansions n the human lineage^29^ These analyses successfully identified an ancestral Catarrhini duplication event, followed by a great ape-specific duplication featuring a lineage-specific 5’ ERV retrotransposon associated with salivary expression^27^ (**Figure 2**). In addition, recent assemblies available for these species suggested lineage-specific duplications within rhesus macaque and olive baboon genomes that were undetected in earlier work.

**Figure 2.**
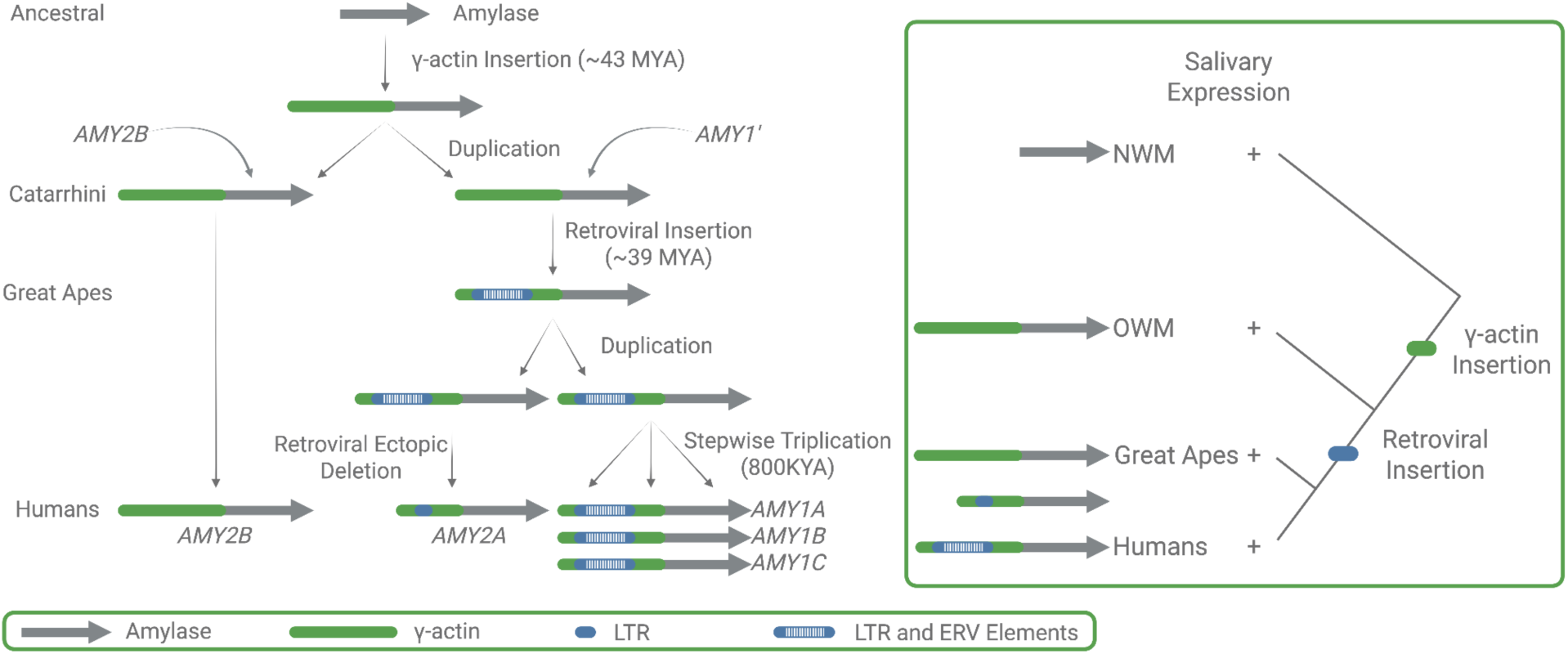
Evolutionary origins of the amylase locus in primates. Model of amylase locus evolution across primates adjusted from Samuelson et al. 1990^27^ The ancestra primate genome contained a single-copy amylase gene (*AMY2B*). n the Catarrhini lineage, a γ-actin insertion occurred upstream of *AMY2B* (∼43 MYA), followed by duplication of the γ-actin-*AMY2B* segment, generating a second gene (AMY1′). n great apes, a retroviral (ERV) insertion occurred (∼39 MYA), followed by retroviral ectopic deletion in the human lineage, giving rise to *AMY2A* In humans, a stepwise triplication of the salivary-expressed amylase gene (*AMY1*) generated *AMY1A AMY1B* and *AMY1C* The right panel summarizes the inferred regulatory and structural events, showing independent gains of salivary expression in New World monkeys, Old World monkeys and great apes, with subfunctionalization of *AMY1* and *AMY2A* in humans.

Our findings provide phylogenetic evidence indicating that the primate ancestor possessed a single amylase gene, orthologous to human *AMY2B* (**Figure 1A**). In the Catarrhini ancestor, after the divergence of new world monkeys, an insertion of the 3’ untranslated region of a γ-actin pseudogene insertion occurred 5’ upstream of the ancestral amylase gene followed by the duplication of the γ-actin-*AMY2B* segment, thereby generating a new amylase copy (*AMY1’)* (**Figure 2**). Later in the great ape lineage, an endogenous retrovirus (ERV) was inserted into the γ-actin region flanking *AMY1’* This combined γ-actin–ERV–*AMY1’* segment duplicated into *AMY1* and the precursor of *AMY2A* in great apes. Eventually, the progenitor of *AMY2A* underwent an ectopic deletion of a portion of the ERV element, leading to its current structure as described further along with mutational mechanisms (**Figure 2**). Our observations within the great ape lineage builds upon previous findings reported by Meisler & Ting^29^ but provides a more granular picture. The observation that *AMY1’* represents the ancestral copy leading to great ape *AMY2A* and *AMY1* is remarkable because these genes have distinct functions with specialized expression in pancreas and salivary glands, respectively.

### Lineage-specific duplications in rhesus macaques

In Old World monkeys, we found that baboons and rhesus macaques retain both the ancestral *AMY2B* gene a well as the derived *AMY1’* gene, which originated in the Catarrhini ancestor (**Figure 2A**). Additionally, we identified one lineage-specific duplicate in macaques and two additional, species-specific duplicates in baboons (**Figure 1A**). To better understand these events, we assessed the gene orthologies of these duplicates and resolved the duplication breakpoints relative to the ancestral Catarrhini and great ape haplotypes (**Figure S3**).

The novel lineage-specific duplication in rhesus macaques, which w termed *AMYm* is located between *AMY2B* and *AMY1’* (**Figure 3A**). Sequence comparisons indicated that *AMYm* exhibits the highest similarity to *AMY2B* suggesting its origin from the ancestral *AMY2B* gene. *AMY1’* in rhesus macaques o the other hand, shares orthology with great apes’ *AMY2A* and *AMY1* although without clear one-to-one orthology with either gene, suggesting that *AMY1’* represents the ancestral state from which *AMY1* and *AMY2A* evolved.

**Figure 3.**
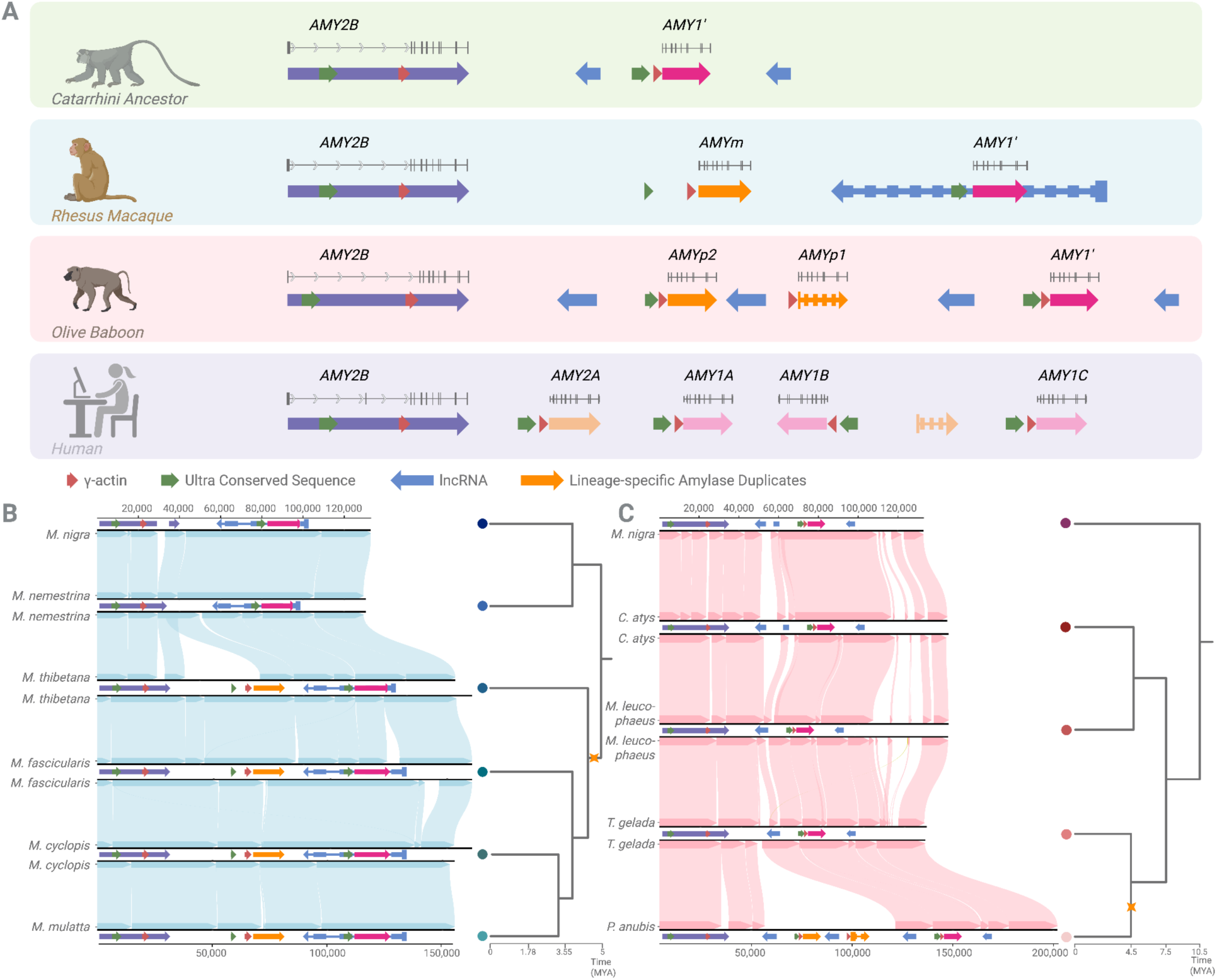
Independent duplication events and structural evolution of the amylase locus in Old World monkeys. (A) Schematic reconstruction of the amylase locus in the Catarrhini ancestor, rhesus macaque (*Macaca mulatta*), olive baboon (*Papio anubis*), and human reference genome (which is also shown to be the ancestral human arrangement^4^ (hg38). The Catarrhini ancestor contains two genes: *AMY2B* and *AMY1′* the latter derived from a duplication of *AMY2B* n rhesus macaques (*Macaca mulatta*), the locus includes *AMY2B AMYm* and *AMY1′* n olive baboons (*Papio anubis*), the locus consists of *AMY2B AMYp2* and *AMYp1* and *AMY1′ AMYp1* s annotated as a pseudogene in NCBI, and is indicated with a dashed outline. *AMYp1* and AMYp2 arose from independent duplication events with distinct breakpoints from those of *AMYm* In the human reference genome, the locus contains *AMY2B AMY2A* and three *AMY1* paralogs, *AMY1A AMY1B* and *AMY1C* The *AMY2B* genes are one-to-one orthologs acros species. AMY1′ in macaques, baboons and the Catarrhini ancestor is orthologous among Old World monkeys and represents the ancestral precursor to human *AMY2A* and *AMY1* genes. (B) Synteny analysis and phylogenetic context of the rhesus macaque-specific duplication. *AMYm* (orange) is located between *AMY2B* (purple) and AMY1′ (pink), and is present only in the fascicularis and sinica macaque groups, but absent in the silenus group. This distribution dates the duplication to approximately 4.5-5 million years ago. The gene structure of *AMYm* is identical for coding sequence to *AMY2B* but differs on the 5’ untranslated region. Breakpoint analysis indicates a nonallelic homologous recombination (NAHR) mechanism. (C) Synteny comparison across Papionini species reveals two independent NAHR-mediated duplication events in olive baboons (*Papio anubis*). The first duplication generated *AMYp1* (orange), and the second produced *AMYp2* (orange); both events must have occurred after the complete split from Guinea baboon (*Papio papio*), within the ∼1.85-million-year window inferred for Papionini divergence. Because of local misassembly in the amylase locus in Guinea baboon (*Papio papio*), we carried out the synteny analysis using closely related *Theropithecus gelada* and *Mandrillus leucophaeus* which are members of the Papionini group, supporting this reconstruction. Species are indicated by Latin names; the corresponding common names and clade memberships are listed in the **Table S5**

By comparing the ancestral Catarrhini haplotype (*i.e.* two-copy haplotype in **Figure 3A**) (see **Methods**) with the rhesus macaque (*Macaca mulatta*) haplotype, we resolved the breakpoints of the duplicated sequences (**Figure 3B**). We identified non-allelic homologous recombination (NAHR) as the primary mechanism driving the segmental duplications. NAHR, in this case, is characterized by two key signatures: first, *AMYm* is flanked by segmental duplications, *AMY2B* at the 5’ end and *AMY1’* at the 3’ end; and second, the duplicated segment containing *AMYm* exhibits mosaic characteristics derived from both flanking regions (**Figure S3**). Additionally, the duplication breakpoints overlap with the γ-actin insertion element, suggesting a possible role of this element in having facilitated the duplication events via NAHR. Among macaques, we were further able to determine that the duplication is specific to *sinica* and *fascicularis* groups and must have occurred after the divergence of the *silenus* group (**Figure 3B**), dating this duplication to the relatively tight window of 4.5 to 5 million years ago based o the previously published phylogenetic dating of this clade^34^ Based o these findings, we constructed what we believe to be the most plausible model that explains the mutational mechanism underlying this macaque-specific duplication event (**Figure S4**).

### Species-specific duplications in olive baboons

We conducted a similar comparative analysis of the olive baboon amylase locus to identify shared and lineage-specific amylase gene duplications and the mechanisms through which they arise Specifically, we investigated the structural configuration of the amylase locus by comparing the olive baboon haplotype carrying four gene copies to the ancestral Catarrhini haplotype. We identified two lineage-specific amylase copies in *Papio anubis* flanked by *AMY2B* on the 5’ and *AMY1’* o the 3’ (**Figure 3C**), which we termed *AMYp1* and *AMYp2*

These two novel genes were absent from other Old World monkey genomes, consistent with independent duplication events specific to the *Papio* lineage. To infer the sequential order of these duplications, we first decomposed the olive baboon amylase locus into four duplicated segments (see **Methods**). These analyses showed that the segment harboring *AMYp1* is a mosaic of the ancestral *AMY2B* and *AMY1’*-containing blocks, whereas the segment carrying *AMYp2* is nearly identical to the *AMYp1* segment, consistent with a second NAHR event duplicating a pre-existing *AMYp1*-containing block. An initial inspection of the Guinea baboon (*Papio papio*) amylase locus revealed six identical tandem amylase segments in addition to the ancestral *AMY2B* and *AMY1’* (see **Methods** and **Figure S5**), thus we treated these redundant copies as unreliable for ordering the duplication events. Instead, we relied on the internal segmental architecture of the olive baboon locus, together with competitive mapping among the four olive baboon *AMY* paralogs, to establish that *AMYp1* represents the first duplication derived from the ancestral *AMY2B-AMY1’* configuration, whereas *AMYp2* arose later as a second, likewise olive baboon-specific duplication, both highlighted in orange in **Figure 3C**

Based on the data sources available to us and our findings outlined above, the most parsimonious mutational model for these olive baboon-specific duplications to be as follows: The ancestral haplotype, containing *AMY2B* and *AMY1′* underwent a initial non-allelic crossover between these two genes, giving rise to the novel *AMYp1* copy. Subsequently, haplotypes carrying *AMY2B AMYp1* and *AMY1′* experienced a second recombination event, this time between *AMY2B* and *AMYp1* resulting in the emergence of the second novel copy, *AMYp2* which is ow positioned between *AMY2B* and *AMYp1* in the extant olive baboon haplotype (**Figure 3C**). Collectively, our reconstructions indicate that *AMYp1* predates *AMYp2* and that both duplications arose after the complete split from Guinea baboons (*Papio papio*). The divergence between olive and Guinea baboons has been dated to approximately 1.85 million years ago based on previously published estimates^35,36^ indicating that both *AMYp1* and *AMYp2* are younger than this split and originated within the olive baboon lineage.

Taken together, our findings highlight lineage-specific duplications in rhesus macaques and olive baboons that occurred independently of one another through NAHR and within approximately 10 million years of evolutionary divergence. Two major questions remain, which we address in the next two sections. First, what was the initial mutational driver of these duplications? Second, what could be the adaptive and functional impact of these lineage-specific duplications?

### Mechanistic inference of NAHR

Our inference of NAHR is based on multiple convergent lines of evidence assessed against the established diagnostic criteria for NAHR, MMBIR and NHEJ^37–39^ First, each lineage-specific duplicate (*AMYm* in macaques; *AMYp1* and *AMYp2* in baboons) is a chimeric copy whose 5′ and 3′ blocks derive from different flanking paralogues, with a single transition point, the hallmark of a strand exchange between misaligned paralogous sequences (**Figure S3**). Second, pairwise alignments reveal that the flanking homologous tracts far exceed the ∼300-500 bp minimal efficient processing segment (MEPS) established for meiotic NAHR^40^ providing ample substrate for homologous recombination. Third, the recurrence of independent events at the same flanking segments across macaques, baboons, bonobos and great apes is itself diagnostic; NAHR characteristically produces clustered, recurrent breakpoints mediated by a fixed pair of segmental duplications, whereas replication-based mechanisms (MMBIR) generate non-recurrent rearrangements with unique breakpoint footprints^37,39^ Finally, we observe neither the complex junction architecture, embedded triplications, inversions, or 2-5 bp microhomology-supported junctions, expected under MMBIR nor the short insertions, short duplications at the break junctions or blunt joins without flanking homology that are characteristic of NHEJ^38^ These observations collectively and robustly support NAHR as the main driver of structural variation in the primate amylase locus.

### The abundance of long terminal repeats (LTRs) correlates with gene copy number gains in the amylase locus

The amylase locus is exceptional in that it has evolved rapidly through independent structural rearrangements across the tree of life, including in species such as fruit flies, mice, rats, and dogs^17,22,41,42^ Several studies have linked this extensive variation to dietary adaptations^17,18,22^ As described above, the mutational mechanisms driving additional duplications, following the initial structural changes in the Catarrhini ancestors, have been characterized as non-allelic homologous recombination (NAHR). However, the origins of primary duplications, which arose from a single-copy ancestral haplotype (likely representing the ancestral state in primates), remain poorly understood.

Our previous work^17^ identified lineage-specific insertions of transposable elements coinciding with *AMY* gene duplications across various mammalian taxa. The association between transposable elements (TEs) and structural variation has been explored in multiple contexts, and recent studies have shown that TE-mediated rearrangements can arise through diverse molecular mechanisms and induce structural instability^43,44^ Building on these observations, we hypothesized that TEs elements might contribute to the formation of primary duplications in primates, thereby predisposing the amylase locus to structural instability. To test this hypothesis, we used a standardized primate repeat dataset to annotate transposable elements across 53 primate genomes, thereby minimizing genome-specific biases (**Table S6**; see **Methods** for details).

Our analyses revealed a wide range of transposable element content in the amylase locus across species, from 25.23% in the northern greater galago (*Otolemur garnettii*) to 59.52% in the Bornean orangutan (*Pongo pygmaeus*) (**Figure 4A**). In contrast to our expectations, we found that the amylase locus is generally depleted in transposable elements compared to genome-wide averages; nevertheless, we observe an enrichment in LTRs (**Figure 4B & 4C**). The general depletion is primarily driven by reduced representation of common, active, short retrotransposons such as Alu sequences (**Figure 4B**), suggesting that the locus is not broadly permissive to transposable element retention.

**Figure 4.**
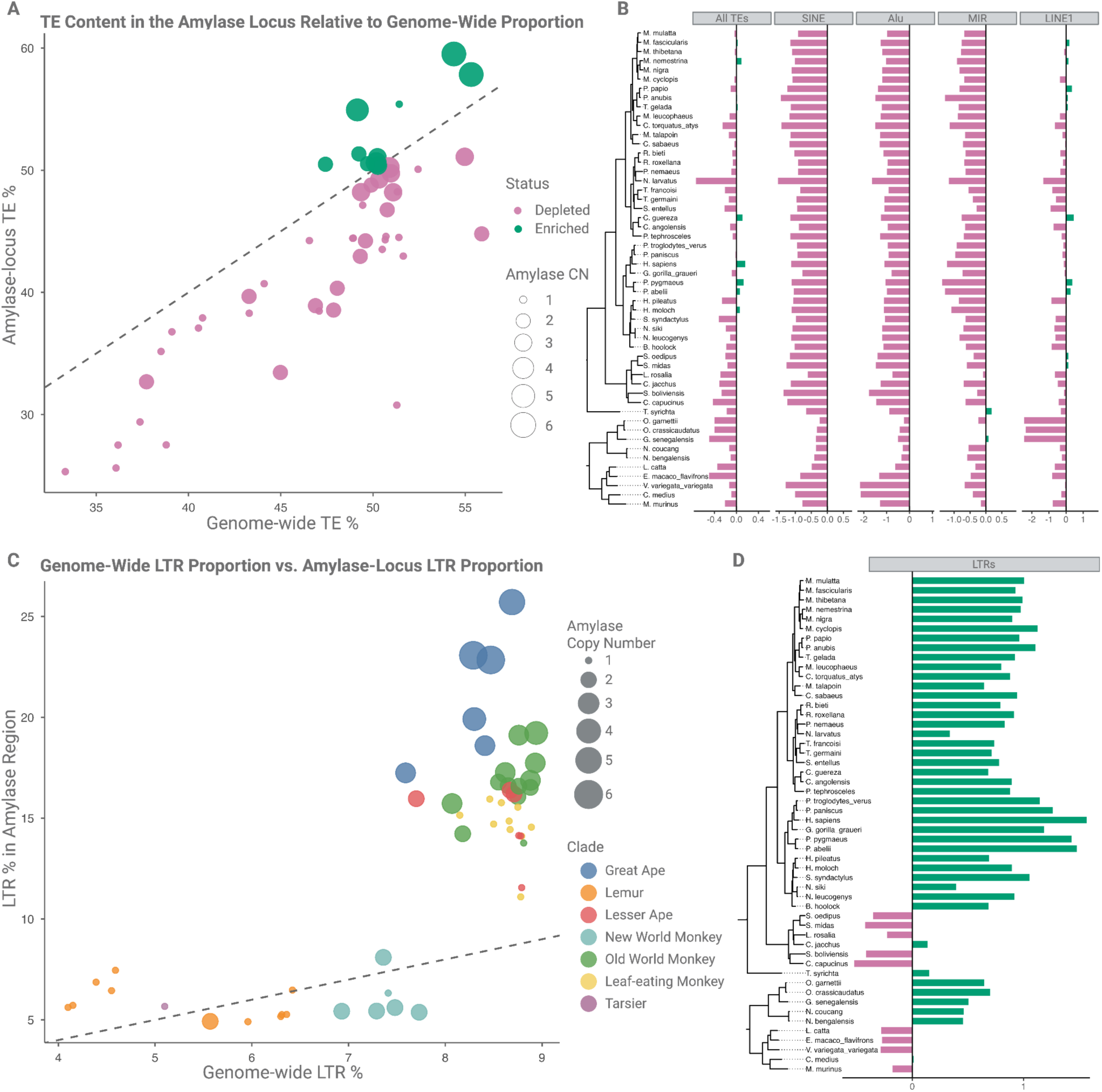
The transposable element landscape in the primate amylase locus. (A) Transposable element (TE) content (given as the total TE in bp/the total length of the locus in bp, see Methods for more) in the amylase locus relative to genome-wide TE proportion across 53 primate species. Each point represents a species, with the size of the circle scaled by the respective amylase copy number and the color indicating enrichment status The dashed line marks the 1: expectation. The majority of species show depletion of TEs in the amylase locus compared to genome-wide levels. (B) TE family-specific enrichment (log_2_ transformed) in the amylase locus across primates for SINEs, Alus, MIRs and LINE1s, alongside total TE content (“All TEs”) (Table S6). Bars represent enrichment (green) or depletion (pink) relative to genome-wide TE representation. Short retrotransposons (e.g. Alus & SINEs) are consistently depleted from the amylase locus, suggesting that this region might have limited retention of active mobile elements across lineages. (C) Relationship between genome-wide and amylase locus-specific LTR content across primate species. Each point represents a species, with the circle size indicating AMY gene copy number and the color representing phylogenetic clade. The dashed line indicates a 1:1 ratio. Most species show an enrichment of LTRs at the amylase locus relative to genome-wide levels. (D) Log_2_-transformed enrichment and depletion of LTR content at the amylase locus across 53 primate species, organized by phylogeny. Bars indicate whether LTR representation in the locus is higher (green) or lower (pink) than genome-wide LTR proportions. Several species with reported amylase gene duplications, such as olive baboon, exhibit pronounced LTR enrichment.

While other common transposable elements were underrepresented within the amylase locus in most primate species, we observed an enrichment of LTR transposons (**Figure 4**, panels **C** and **D**). The proportion of LTR elements in the locus, even after accounting for phylogenetic relatedness, correlates with the number of amylase genes (p 10^-5^ R^2^ 0.62; **Figure S6**). This correlation is most significant within Catarrhini, where the majority of amylase gene gains and losses were observed, supporting the hypothesis that LTRs contribute to structural instability in the amylase locus f indeed LTRs are driving the structural changes, we would expect to find LTRs proximal to the breakpoint junctions, as exemplified by the *AMYp2* duplication in olive baboons (**Figure S7**). We also detected a strong negative correlation between amylase gene copy number and the presence of DNA transposons, particularly *Charlie* and *Tigger* elements (p 1.4e^-6^ and p = 0.0005 respectively; **Figure S8**), further raising the possibility that non-LTR transposons might have limited retention in the locus. Together, these findings point to a transposable element-specific pattern of enrichment and depletion within the amylase locus, with these dynamics showing strong correlations with amylase gene copy number variation across primates.

To further assess the relationship between LTRs and segmental duplications, we identified orthologous LTR insertions across primates, clustered them into orthogroups, and reconstructed ancestral presence/absence states on the primate phylogeny (**Figure S9 & S10**). This analysis revealed elevated LTR gain events on the branch leading to Catarrhini, coinciding with the initial amylase duplication events in this lineage. Importantly, for the LTRs discussed above, the structural evidence indicates that the LTR copies were part of the ancestral segment that was subsequently duplicated by NAHR, rather than secondary insertions into pre-existing duplicates. Although we cannot conclusively demonstrate LTRs as the direct cause of structural rearrangements, we hypothesize that their insertion may contribute not only to structural instability but may also facilitate functional changes in regulatory regions as suggested earlier^29^ Due to their role in regulatory rewiring, their retention may generate long stretches of homologous sequence, which in turn may cause subsequent NAHR events Such dynamics n primates may explain both the recurrence of amylase gene duplications, as well as their functional changes in tissue expression.

### Signals of selection suggest potential functional variation among primate amylase paralogs

While human amylase paralogs in general appear to evolve under negative selection without significant amino acid divergence from each other^4^ recent studies highlighted potentially functional variations among human paralogs^45^ To identify such functional differences and signals of potential positive selection, we aligned the coding sequences of all identified paralogs from olive baboons and rhesus macaques with paralogs from ape genomes.

Our analysis yielded three major insights (**Figure 5A & 5B)** First, we identified a significant positive selection signal on the internal branch leading to baboon and rhesus macaque paralogs compared to great ape paralogs (aBSREL, p = 0.038). Second, we detected a premature stop codon mutation in the ancestral *AMY2B* gene copy within baboons, previously unannotated in the NCBI database (**Figure 5A**). Third, we detected strong evidence for positive selection specifically on *AMYp2* in olive baboons, supported by both aBSREL (p 10^-6^) and RELAX analyses (p 0.0001). Additionally, we identified six codons exhibiting episodic positive selection that likely contribute to this overall selection signal (**Methods; Figure S11**). The concurrent presence of a premature stop codon in *AMY2B* and positive selection in *AMYp2* is remarkable. It has been shown previously that following gene duplications, one of the copies may become pseudogenized and the other one retains the original function^46,47^ Thus, it is plausible here that the newly derived gene (*AMYp2*) functionally compensates for the observed loss of function in the ancestral gene (*AMY2B*).

**Figure 5.**
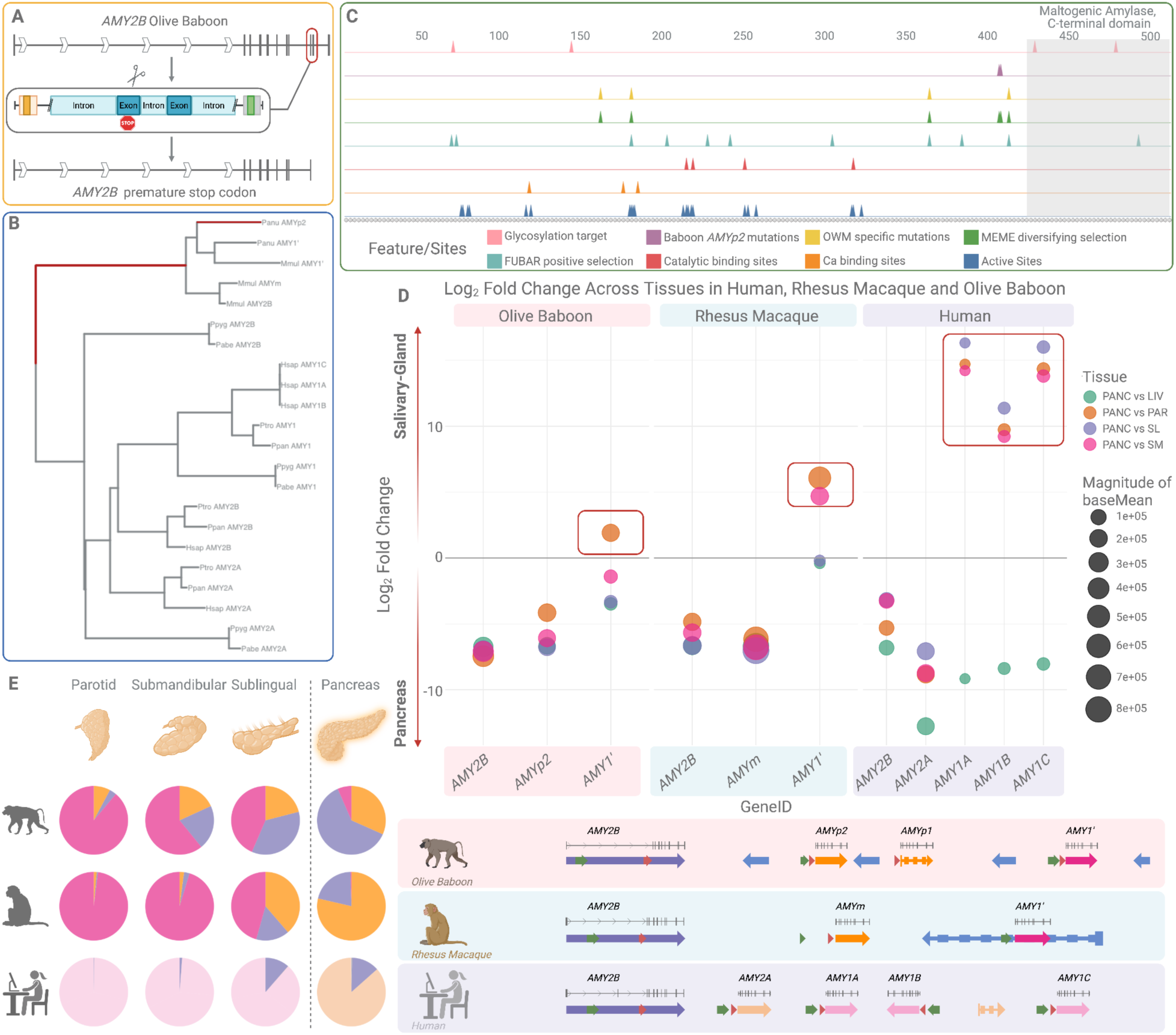
Functional divergence and tissue-specific expression of amylase paralogs in olive baboons, rhesus macaques and humans. (A) Identification of a premature stop codon in *AMY2B* of olive baboon. A schematic of the disrupted gene structure (top) shows the location of the nonsense mutation (red box) within the ninth exon. (B) A maximum likelihood phylogeny of amylase coding sequences from olive baboon, rhesus macaque and great apes, highlighted branches under significant episodic diversifying selection (dark red) affecting the lineage leading to Old World monkey paralogs (aBSREL, p = 0.038). (C) Old world monkey and baboon-specific variants and their overlap with predicted functional sites across the amylase protein. Tracks show the position of catalytic and calcium-binding residues, glycosylation sites, positively selected codons (FUBAR, MEME), Old World monkey-specific mutations and baboon-specific mutations in *AMYp2* A MEME- and FUBAR-identified site (codon position 178) involves a threonine-to-serine substitution overlaps an active site. (D) Differential expression of amylase paralogs across tissues in olive baboons, rhesus macaques and humans. The scatterplot shows log_2_-fold change for each paralog across tissue comparisons (pancreas vs parotid, sublingual or submandibular glands and pancreas vs liver). The size of the circle corresponds to the base mean expression evel. In baboons and macaques, lineage-specific genes (*AMYp2 AMYm*) contribute to salivary expression, unlike in humans where *AMY1* is the dominant/primary salivary paralog. Arrows represent the structure and orientation of amylase paralogs across species, colored by paralog identity: *AMY2B* (purple), *AMY2A* (light orange), *AMY1* (light pink), *AMY1′* (pink), *AMYm AMYp1* and *AMYp2* (orange). (E) Pie charts showing proportional expression of each amylase paralog in salivary glands (parotid, submandibular, sublingual) and pancreas. Expression levels of *AMY1′* in both salivary glands and pancreas in macaques and baboons supports subfunctionalization as the likely evolutionary mechanism for the tissue-specific roles of *AMY1* and *AMY2A* in humans.

Next, we analyzed the functional annotations of primate amylase amino acid sequences, including known glycosylation sites, streptococci-binding motifs, catalytic and active sites, and calcium-binding domains (**Figure 5C** see **Methods**). To evaluate whether the observed amino acid substitutions might affect protein structure or function, we generated AlphaFold2 models for the amylase gene products of each paralog, with a particular focus on the baboon-specific *AMYp2* sequence which showed the strongest signal of positive selection (**Figure S12**). The functional predictions suggest that most of the positively selected substitutions do not overtly disrupt the protein fold, glycosylation motifs, or key catalytic residues. However, subtle impacts on substrate affinity or protein stability cannot be ruled out (**see Table S7** for predicted pathogenic and neutral substitutions). Notably, we identified one strongly selected site (p = 0.01, MEME) in *AMYp2* involving a threonine-to-serine substitution at the position 178 predicted as an active site by NLM’s conserved domain database^48^ which may reflect fine-tuned functional divergence in the olive baboon lineage. Collectively, our findings suggest that, unlike reported for humans^4^ Old World monkey amylase gene duplications involve significant amino acid diversification, indicative of potential neofunctionalization.

### Reconstructing the evolution of salivary gland-specific expression

To investigate the expression patterns of lineage-specific amylase gene paralogs, we generated transcriptome data from parotid, sublingual, and submandibular salivary gland, as well as liver and pancreas tissues from six rhesus macaques and five olive baboons (**Table S8**). We also leveraged our previously published transcriptomic data from the corresponding human salivary gland tissues^49^ and added data for pancreas and liver tissues from the GTEx database. Taking advantage of sequence differences among paralogs within each species, we used Kallisto^50^ to quantify transcript abundance of each gene as it offers high sensitivity and accuracy for distinguishing among closely related gene copies^50^ (see **Methods** for more detailed discussion). These findings led us to form several hypotheses to explain the regulation of amylase duplicates.

The first clear pattern we observed in both baboons and rhesus macaques is that, similar to humans, the last gene (at the 3’ end) in the amylase cluster consistently shows elevated expression (parotid vs pancreas log_2_FC: rhesus *AMY1’*=6.08, baboon *AMY1’*=1.91) in salivary tissues relative to the other paralogs (**Figure 5D**). However, in contrast to humans, where *AMY1* is exclusively expressed in saliva, this gene in baboons and rhesus macaques retains expression (pancreas vs liver log_2_FC: rhesus *AMY1′*=0.40, baboon *AMY1’*=3.48) in the pancreas and liver, indicating broad expression patterns (**Figure 5E)** Additionally, the relative contribution of each amylase paralog to total salivary expression differs among species (**Figure 5D**). In humans, *AMY1* accounts for nearly all salivary gland expression (parotid vs pancreas log_2_FC: *AMY1A*=14.71, *AMY1B*=9.74, *AMY1C*=14.35). The lineage-specific duplications *AMYm* and *AMYp2* also contribute to the overall salivary expression in rhesus macaques and olive baboons, respectively.

Given that *AMY1’* in Old World monkeys represents the ancestral gene from which the great ape *AMY1* and *AMY2A* genes evolved (**Figure 2A**), it offers a valuable framework for investigating how tissue-specific expression arose in these newly duplicated genes. In humans, *AMY1* is expressed exclusively in salivary glands, while *AMY2A* is expressed only in the pancreas (**Figure 5E**). Our transcriptomic analyses in rhesus macaques and baboons shows that *AMY1’* is expressed in both pancreas and salivary glands, likely reflecting the ancestral state for Catarrhini (**Figure 5D & 5E**). Based on these observations, the most parsimonious explanation is that the ancestral *AMY1’* gene had already acquired expression in both the pancreas and salivary glands, particularly the parotid gland, prior to the divergence of great apes. Following duplication in the great ape lineage, subfunctionalization occurred: *AMY1* retained salivary gland expression, while *AMY2A* lost salivary expression and became restricted to the pancreas. This shift may have been facilitated by an ERV insertion, as previously proposed^27,29^ Together, these findings support a subfunctionalization model for the great ape *AMY1* and *AMY2A* following their divergence from *AMY1’*

### Multifactorial regulation of tissue-specific expression of amylase paralogs

To investigate how regulatory elements contributed to the evolution of amylase gene expression in primates, we examined transcription factor binding sites across paralogs and species. Using *in silico* predictions of transcription factor binding sites and promoter regions for amylase paralogs in rhesus macaques olive baboons, and humans, we identified 262 distinct binding sites (consisting of 108 unique TFBS) across the promoters of the nine annotated amylase gene paralogs analyzed (**Table S9**).

A simplistic model of tissue-specific expression assumes that the presence of a specific transcription factor binding site determines tissue specificity. If this were the case, we would expect the transcription factor binding motifs among primate amylase paralogs to cluster according to their expression in either salivary glands or pancreas However, we found no such pattern: paralogs with known tissue-specific expression did not show consistent enrichment of salivary- or pancreas-biased motifs (**Figure 6**). Instead, our results suggest a partial conservation in promoter binding site composition among primate paralogs, where all three rhesus macaque paralogs (*AMY2B AMYm* and *AMY1’*) share a similar binding site profile with olive baboon *AMY1’* and *AMYp2* In contrast, the promoter regions of the pseudogenized olive baboon *AMY2B* and the human paralogs (*AMY2B AMY2A* and *AMY1s*) exhibit a distinct transcription factor binding sequence composition, consistent with the lineage-specific evolution of these gene copies (**Figure 2A**).

**Figure 6.**
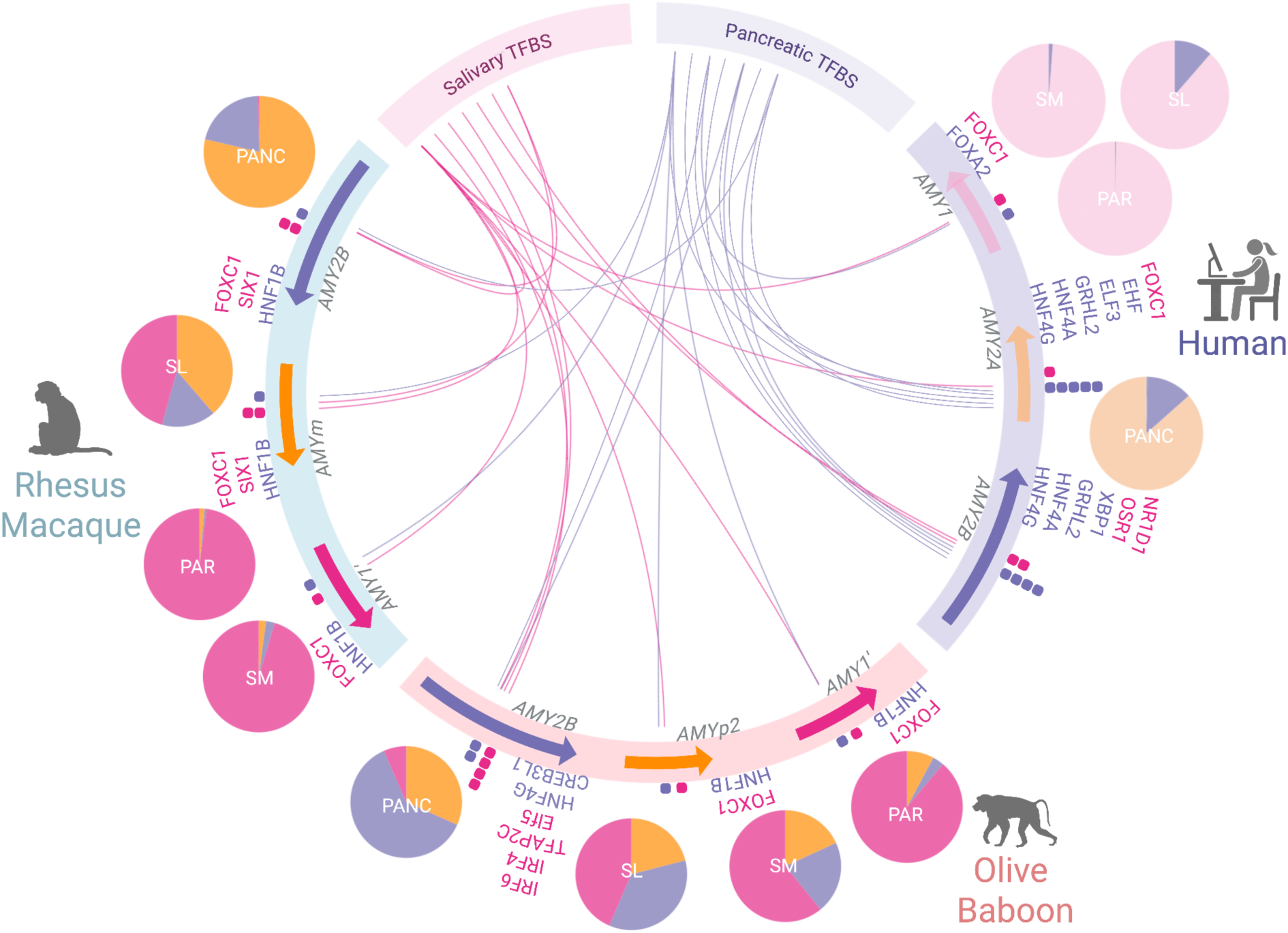
Predicted transcription factor binding sites (TFBS) associated with salivary-gland and pancreatic expression across amylase gene paralogs. Circos plot showing in silico predictions of salivary (pink) and pancreatic (purple) TFBS across promoter regions (∼170 bp upstream of the start codon and annotated promoter) of amylase paralogs in rhesus macaques (*Macaca mulatta*), olive baboons (*Papio anubis*) and humans. Colored arrows represent gene orientation. Dots adjacent to each gene denote the presence of TFBS (pink for salivary, purple for pancreatic). Ribbon links connect paralogs to TFBS categories. Pie charts summarizing relative gene expression across tissues: PAR (parotid), SM (submandibular), SL (sublingual), and PANC (pancreas). The color of each slice corresponds to the proportion of total expression of the given gene in each tissue, following the sam color scheme used for the gene annotation arrows All three rhesus macaque (*Macaca mulatta*) paralogs (*AMY2B AMYm AMY1′*) share similar TFBS profiles with olive baboons (*Papio anubis*) *AMY1′* and *AMYp2* In contrast, *AMY2B* in olive baboon and human exhibit distinct TFBS composition. The salivary gland-biased transcription factor FOXC1 is present in most paralogs, but absent from human and olive baboon *AMY2B*, both of which lack salivary expression.

FOXC1 is a key transcription factor involved in salivary gland development and expression^51^ Notably, we found that all primate *AMY* paralogs, regardless of whether they are salivary gland- or pancreas-biased, contain FOXC binding sites, with the exception of human *AMY2B* and the pseudogenized olive baboon *AMY2B* This observation suggests that the presence of a salivary-biased regulatory motif alone is insufficient to dictate tissue-specific expression. Instead, expression patterns of *AMY* genes may be further shaped by context-dependent factors such as chromatin accessibility, competitive binding, or the presence of co-regulators. For example, despite containing a FOXC1 binding site, human *AMY2A* is primarily expressed in the pancreas, highlighting the complexity of regulatory control at this locus.

These findings suggest that the presence or absence of salivary gland or pancreas-biased transcription factor binding sites alone does not fully account for the observed tissue-specific expression patterns, highlighting the potentially combinatorial nature of transcriptional regulation, an observation consistent with previous findings from other tissue-specific systems^49^

## DISCUSSION

In this study, we used the amylase gene locus as a model to investigate within the primate lineage how structurally complex loci contribute to molecular convergence Convergent evolution, where similar traits arise independently across lineages, is a hallmark of adaptive evolution and highlights non-random patterns shaped by natural selection. Structurally complex regions have recently emerged as key players in this process, and it is thought that independent gene duplications might play an important role in this context. Focusing on the amylase locus, which is one of the most structurally dynamic regions in the human genome^4^ we expanded prior work in humans to include the broader primate phylogeny. We identified multiple lineage-specific amylase gene duplications, including previously uncharacterized expansions in bonobos, orangutans, lemurs, and New World monkeys. This comprehensive analysis of amylase evolution enabled us to dissect the interplay between mutational mechanisms, positive selection, and regulatory divergence underlying functional convergence.

Our findings revealed that NAHR-mediated rearrangements in baboons, macaques, and humans occurred through distinct breakpoints, underscoring the recurrent nature of duplication events. Given that NAHR relies on existing homologous sequences, we asked how primary duplications arise in the first place. We observed a strong correlation between amylase copy number and LTR abundance across species, consistent with the hypothesis that lineage-specific transposable element insertions may contribute to the homology required for initiating structural instability. While we cannot exclude a bidirectional relationship, in which segmental duplications create a permissive genomic context for additional TE accumulation, the synteny-based evidence is consistent with LTRs providing additional stretches of shared sequence that facilitate NAHR This provides preliminary evidence for a broader hypothesis, that transposable elements may be a plausible contributor in priming structurally complex loci for recurrent evolution, a compelling direction for future studies across mammalian genomes.

Beyond structural variation, we detected signatures of positive selection at specific codons, suggesting functional divergence among amylase paralogs in a species-specific manner. Such changes may be of functional relevance because they can alter the conformation of the protein and change or create motifs governing posttranslational modifications. Evidence supports that human AMY paralogs differ in glycosylation potential and may influence oral microbiota composition^52^ The interplay of nucleotide-level and structural variation lays the groundwork for future population-level studies in closely related species with differing diets or pathogen pressures, offering a powerful framework to investigate the adaptive relevance of *AMY* variation.

By integrating genomic variation, gene expression, and transcription factor motif analyses, we show that *AMY* paralogs have independently evolved tissue-specific expression, particularly in salivary glands. Rather than relying on distinct, tissue-specific transcription factors, expression biases appear to result from varying combinations of binding motifs among paralogs and across species. This regulatory rewiring is linked to structural rearrangements at the locus, including duplications, inversions, and deletions, that reshape the genomic context of regulatory elements. Our findings, thus, highlight the likely importance of distal regulatory elements beyond core promoters in mediating tissue-specific expression. Future functional assays such as ATAC-seq, ChIP-seq, or Fiber-seq will be essential to fully resolve the evolving regulatory landscape of the amylase locus.

A vivid example of the co-evolution of gene duplication and regulatory rewiring is seen in the great ape *AMY2A* and *AMY1* genes Our results show that these paralogs arose from an ancestral gene (*AMY1’*) with dual expression in the pancreas and salivary glands. Following duplication, *AMY2A* and *AMY1* subfunctionalized into largely pancreas- and salivary gland-specific expression, respectively. This instance of regulatory and functional co-divergence exemplifies the distinct evolutionary innovation at the primate amylase locus, ranging from fixed duplications (as in macaques and baboons), to extensive intraspecific variation (as in humans), and even gene loss (in leaf-eating monkeys).

Notably, we found no species lacking the amylase gene entirely, suggesting that while the locus tolerates considerable structural and regulatory flexibility, it remains under consistent functional constraint. This balance, between mutational plasticity and adaptive necessity, places the amylase locus in what Ponting^53^ has termed the “evolutionary twilight zone”. It is within this zone that convergence has repeatedly emerged during primate evolution, driven by distinct molecular mechanisms acting on a structurally complex genomic landscape.

We propose that structurally complex loci across mammalian genomes are hotspots of molecular convergence, harboring exceptional evolutionary potential. Their structural complexity not only promotes gene copy number expansion and contraction but also enables regulatory innovation, fueling expression divergence and functional diversification of duplicated genes.

### Limitations of the study

Several limitations of this study should be noted. First, our analyses rely on a limited number of genome assemblies (at most 4) per species. While parsimony-based reconstruction of copy number evolution shows clear phylogenetic concordance and we confirmed structural configurations on multiple assemblies for 16 species, we cannot exclude within-species copy number polymorphism in the amylase locus. Population-level surveys, analogous to what has been achieved for humans^3,4,20^ will be essential to characterize the full extent of structural variation at this locus across primates.

Second, genome assembly quality can influence TE annotation, particularly in repetitive regions. We showed that LTR annotations are broadly consistent between short-read and long-read assemblies for the same species and that the significant correlation between assembly N50 and genome-wide TE content is primarily driven by satellite sequences rather than LTRs Further, we identified one case in the Guinea baboon (*Papio papio*) in which local misassembly of the amylase locus substantially affected TE estimates. We resolved this issue by performing targeted local reassembly using the original long reads and re-annotating transposable elements on the corrected locus. Despite these efforts, we cannot entirely rule out similar, undetected local assembly errors in other species, though no additional outliers were observed.

Third, the causal direction between LTR enrichment and amylase gene copy number gains remains ambiguous. In SD-rich regions, NAHR can both expand copy number and create a genomic context permissive to additional TE accumulation, making it difficult to disentangle cause from consequence. Our synteny-based evidence indicates that LTR insertions were present in the ancestral segment before duplication, consistent with a seeding role; however, we acknowledge that TE insertions and segmental duplications likely influence each other bidirectionally at this locus.

Finally, our regulatory analysis is based on *in silico* TFBS prediction and does not directly measure chromatin accessibility or transcription factor occupancy. The finding that FOXC1 motifs are present in most *AMY* promoters regardless of tissue-specific expression underscores the insufficiency of motif-based prediction alone and highlights the need for functional assays (e.g., ATAC-seq, ChIP-seq, Fiber-seq) to resolve the regulatory architecture of the amylase locus.

## RESOURCE AVAILABILITY

### Lead contact

Requests for further information and resources should be directed to and will be fulfilled by the lead contact, Omer Gokcumen.

### Materials availability

All unique materials generated in this study are available from the lead contact with appropriate institutional approvals.

### Data and code availability

Bulk RNA-seq expression data (TPM, GTEx v8) are available from the GTEx open-access portal (https://gtexportal.org/home/downloads/adult-gtex/bulk_tissue_expression). RNA-seq datasets for olive baboon (*Papio anubis*) and rhesus macaque (*Macaca mulatta*) have been deposited at GEO under accession numbers GSE305241 and GSE305255, respectively. All other data reported in this paper are available in the main text and supplementary information.

Al original analysis scripts and input files for downstream analyses and visualization have been deposited at Zenodo and are publicly available at https://doi.org/10.5281/zenodo.16809248 and https://doi.org/10.5281/zenodo.18689074 This paper does not report any additional original code.

Any additional information required to reanalyze the data reported in this paper is available from the lead contact upon request

## ACKNOWLEDGMENTS

We thank Leo Speidel, and Luane Landau for carefully reading this manuscript.

## AUTHOR CONTRIBUTIONS

Conceptualization, C.K. and O.G.; methodology, C.K; Investigation, C.K.; C.K. conducted the bioinformatics analyses; P.P. conducted the ddPCR assay for within-species variation analysis; writing—original draft, C.K. and O.G.; writing—review & editing, C.K., S.R., O.G.; funding acquisition, S.R. and O.G.; resources, O.G.; supervision, O.G.

## DECLARATION OF INTERESTS

The authors declare no competing interests.

## DECLARATION OF GENERATIVE AI AND AI-ASSISTED TECHNOLOGIES

During the preparation of this manuscript, the authors used ChatGPT for language and grammar checks. After using this tool or service, the authors reviewed and edited the content as needed and take full responsibility for the content of the publication.

## SUPPLEMENTAL INFORMATION

**Document S1. Glossary & Figures S1–S14 (pdf file) Document S2. Tables S1–S18 (Excel file)**

## EXPERIMENTAL MODEL AND STUDY PARTICIPANT DETAILS

Macaque and Baboon salivary glands samples were obtained from Texas Biomedical Institute through their established ACUC protocols and by expert veterinary staff. The samples were obtained opportunistically during planned euthanasia from animals that are 7 to 18 years old (**Table S8)**

## METHOD DETAILS

### Primate Genome Assemblies and Locus Extraction

To capture the phylogenetic diversity of the amylase locus across primates, we analyzed 244 high-quality genome assemblies representing 222 species (**Table S1**). To assess copy number variation and structural organization, we first delineated the locus using two flanking, non copy-number-variable anchor sequences: one upstream (5′) of the *AMY2B* gene corresponding to the *RNPC3* genic region, and one downstream (3′) of the *AMY1C* gene. Both sequences were derived from the human GRCh38 reference genome and used as queries in blastn (v. 2.14.1+)^54^ searches against each assembly.

We automated this analysis using a custom SLURM-based Bash pipeline. The script identified assemblies where both anchors mapped to a single contig or scaffold, extracted the intervening locus using samtools faidx (v. 1.22)^55^ and generated reverse complements with seqtk (v. 1.5-r133; https://github.com/lh3/seqtk) for loci located on the reverse strand. Only assemblies in which the full amylase locus was contained within a single contig were retained for downstream analyses; loci fragmented acros multiple scaffolds were excluded but recorded. The queried sequences and the SLURM-based Bash pipeline have been deposited in Zenodo; see Data Availability.

### Synteny Analyses and Gene Annotation

To characterize the structural composition and assess synteny of the amylase locus across primates, we analyzed the 70 assemblies in which the complete locus was successfully extracted as a single contiguous sequence. Each locus was aligned to the human H1^a^.1 haplotype, which carries a single *AMY1* gene and represents the ancestral configuration in great apes, using NUCmer (v. 3.1)^56^ and LAST^57^ Dotplots were then generated using mummerplot (v. 3.5)^56^ and visually inspected to identify structural rearrangements relative to the human reference.

To further investigate patterns of structural evolution both within and between genera, we performed additional pairwise alignments among species of the same genus, as well as representative comparisons across genera These alignments were again generated with NUCmer (v. 3.1) and visualized using dotplots and miropeats-style plots. For the latter, we used custom scripts to convert NUCmer coordinate files into PAF format and visualized the alignments using the R package SVbyEye (v. 0.99.0) ^58^

To annotate amylase gene copies across primates, we employed a multi-step approach combining reference annotations, CDS mapping, and manual curation. For each genome assembly with a contiguous extracted amylase locus, we first retrieved available NCBI gene annotations for that species and used exonerate (v. 2.4.0)^59^ to map the corresponding coding sequences (CDS) to the extracted regions. In assemblies lacking gene annotations, we utilized CDS from closely related sister species and mapped them to the respective genomes using the same exonerate-based pipeline, which is well-suited for aligning sequences in the presence of sequence divergence and detecting partial exons.

To identify putative orthologs and resolve paralogous relationships, we implemented a recursive reciprocal BLAST approach. Putative amylase gene models from each species were queried against one another using blastn (v. 2.14.1+) and reciprocal searches were used to establish one-to-one or one-to-many orthology/paralogy relationships. In a one-to-one orthology relationship, reciprocal BLAST searches yield the same gene pair as the top hit in both directions e.g gene X from species 1 identifies gene Y from species 2, and gene Y reciprocally returns gene X. By contrast, a one-to-many pattern arises when duplication has occurred in only one lineage, and this can appear in two complementary ways. If species 2 experienced the duplication, single gene in species 1 matches several genes in species 2, and each of those paralogs reciprocally lists the original query gene as its best hit. Conversely, if the duplication occurred in species 1, multiple paralogs in that species all identify the same ortholog in species 2 as their top hit, while the lone gene in species 2 reciprocally aligns most strongly to just one, commonly the most conserved, of the copies in species 1. Either configuration reveals a one-to-many orthology generated by a lineage-specific duplication event This approach allowed s to confidently distinguish ancestral copies (e.g. *AMY2B*) from lineage-specific duplications.

Lastly, in olive baboons (*Papio anubis*), we detected a premature stop codon within the *AMY2B* gene suggesting likely pseudogenization. This mutation is absent from the NCBI annotation and was not present in the CDS of Guinea baboon (*Papio papio)* but was consistently identified in both haploid assemblies of olive baboons.

### Transposable Element Annotation and Quantification

All 53 primate assemblies that yielded a contiguous amylase locus were analyzed using RepeatMasker (v. 4.1.5; http://repeatmasker.org) with the -species primates flag and default parameters. This approach applies a uniform primate-specific repeat library to every genome, avoiding the inconsistencies that would arise from mixing lineage-specific libraries. However, this choice can systematically underestimate species-restricted transposable element (TE) families absent from the reference library, an acknowledged limitation of our approach. RepeatMasker output files (.tbl and .out) were parsed to calculate genome-wide TE content for each assembly. Assembly quality introduces additional bias to these estimates, as chromosome-level assemblies often recover TE-rich centromeric and telomeric regions that incomplete or nearly-complete assemblies commonly miss, thereby inflating total TE proportions.

To analyze TE content within the amylase locus, we applied RepeatMasker separately to the extracted amylase sequences from each of the 53 species. These sequences were bounded by non copy-number-variant *RNPC3* 5′ and *AMY1C* 3′ flanks (detailed in the previous section) and restricted to loci assembled as single, gap-free contigs/scaffolds, as a prerequisite to avoid potential segmental duplication collapse. Masking isolated locus sequences rather than extracting annotations from whole-genome GFF3 files, prevented coordinate errors and ensured that every species was compared across identical boundaries.

TE abundance was quantified as the proportion of masked bases (bp masked / region length) for both genome-wide and locus-specific analyses. This normalization accounts for length variation introduced by gene duplications. All downstream statistical analyses and visualizations were conducted in R (v. 4.3.3). For each species, we compared the percentage of masked sequence in the amylase locus to its genome-wide percentage. For each TE family, we calculated log2-enrichment, defined as log2(% in locus)/(% genome-wide). This metric was then merged with amylase copy number estimates and clade assignments. Enrichment or depletion was assessed with a two-sided Wilcoxon signed-rank test across the 53 species in which the locus was contiguous; family-specific analyses (e.g. Alu, LTR, DNA transposons) were performed analogously. P-values were adjusted for multiple testing using the Benjamini-Hochberg procedure, considering FDR<0.05 significant.

We assessed the potential confounding effect of assembly contiguity (scaffold N50/Assembly N50) on TE content estimates Assembly N50 showed a strong positive correlation with genome-wide TE content (Spearman ρ=0.554, P<0.001; n=53). However, this relationship was primarily driven by assembly length. After controlling for total assembly length, the association between N50 and TE content remained significant but was substantially reduced (partial correlation ρ=0.327, P=0.018), indicating that assembly quality has a modest independent effect on TE detection beyond simple genome size effects.

To directly evaluate the impact of sequencing technology on TE estimation, we leveraged two species for which both short-read-based and long-read-based assemblies are available (*Semnopithecus entellus* and *Macaca nemestrina*). For each species, we a RepeatMasker o both assemblies with identical parameters and quantified the genome-wide proportion of sequence annotated as major TE classes (LINEs, SINEs, LTR retrotransposons, DNA transposons, satellites/low-complexity). Across both species, total TE content and the relative contributions of LINEs, SINEs and LTR retrotransposons were highly similar between short-read and long-read assemblies, whereas the long-read assembly in *Macaca nemestrina* recovered a higher fraction of sequence annotated as satellites and related tandem repeats, consistent with improved representation of centromeric and pericentromeric regions in more contiguous genomes (**Table S10 & S11**).

To test whether TE abundance predicts amylase gene copy number while accounting for shared ancestry, we fit phylogenetic generalized least-squares (PGLS) models in R (nlme 3.1 and caper 1.0.1). A ultrametric primate tree pruned to the 53 focal species was obtained from TimeTree (accessed January 2025). The response variable was amylase gene copy number while the predictor was locus-specific TE proportion (total or by family). Pagel’s λ was estimated by maximum likelihood and retained if significantly different from zero Model fit was evaluated by R^2^ and residual diagnostics. All scripts for TE analysis, including those used to reproduce Figure 4, are available in a Zenodo repository (see Data Availability)

### LTR Orthogroup Analysis and Ancestral-State Reconstruction

To assess the evolutionary dynamics of individual LTR insertions at the amylase locus across primates, we identified orthologous insertions using a reciprocal flanking-sequence approach. For each LTR element annotated by RepeatMasker within the extracted amylase locus of each species, we extracted 1kb flanking sequences on both sides and used reciprocal BLAST to identify orthologous insertions across species. Elements sharing orthologous flanking context were clustered into orthogroups. A binary presence/absence matrix (214 orthogroups x 53 species) was then constructed and analyzed on the pruned primate phylogeny.

We computed parsimony steps per orthogroup using phangorn v.2.12^60^ and retained orthogroups requiring ≤10 changes across the tree which included all trees (n=214). For each retained orthogroup, we reconstructed ancestral states using maximum-likelihood discrete-character analysis (ace function in ape v.5.8)^6^ under a equal-rates (ER) model. MAP ancestral states were assigned to internal nodes, and branch-wise gains and losses were tallied by comparing parent-child state pairs across all edges and orthogroups. Results were visualized by painting branch colors proportional to the total number of LTR gains or osses o the primate phylogeny (**Figure S9 & S10**). All scripts are deposited in the Zenodo repository (see Data Availability).

### Structural Variant Inference and Duplication Mechanism Analysis

To investigate the mechanisms underlying the duplication in the amylase locus, we extended our annotation of the region beyond the existing annotations available o NCBI We utilized BISER (v. 1.4)^62^ to identify segmental duplications in the olive baboon amylase locus, analyzing the masked and unmasked genome to capture all duplicated units, irrespective of functional annotation. This approach led to the identification of 43 distinct duplicated segments (**Table S12**). Additionally, we identified four lncRNA sequences within the locus, positioned downstream of each gene with reverse orientations relative to the genes. Using self-alignments, dotplots, and BLAST searches, we partitioned the amylase locus into four distinct segments, each corresponding to a single duplicon (**Figure S13A**). Each segment includes the coding sequence of an amylase gene and extends to the terminal sequence of an adjacent lncRNA. This partition enables a systematic examination of duplicated regions across the locus.

We then compared the four distinct segments to one another using pairwise NUCmer alignments and visualized these comparisons with dotplots. Query coverage and sequence similarity metrics allowed us to infer the order of duplication events A closer examination of the nucleotide sequences further informed us of the duplication mechanisms involved. Namely, the third segment (containing the amylase gene *AMYp1*) appears to be a composite of the fourth segment (encompassing *AMY1’*) and the first segment (encompassing *AMY2B*) (**Figure S13B**). Meanwhile, the second segment is nearly identical to the third, with the only difference being a ∼10 kb deletion within the third segment (**Figure S13C**). Because this interval is bounded by long stretches of Ns in the olive baboon assembly, we cannot determine whether it represents a true biological deletion or a local misassembly/scaffolding artifact, and we therefore do not interpret this feature further. Taken together, the sequence-similarity patterns indicate that two non-allelic homologous recombination (NAHR) events duplicated the ancestral *AMY2B* and *AMY1′* blocks, yielding the present-day amylase locus structure in olive baboons (**Figure S13**).

To delineate precisely the novel duplications in the olive baboon amylase locus, we utilized closely related Papionini and Old World monkey species that still retain the ancestral Catarrhini two-copy configuration (*AMY2B* and *AMY1′*). Continuous contigs spanning the locus were available for gelada (*Theropithecus gelada*), drill (*Mandrillus leucophaeus*), sooty mangabey (*Cercocebus atys*), and crested macaque (*Macaca nigra*). We extracted the orthologous intervals (*RNPC3* 5′ flank to *AMY1C* 3′ flank; see above) and aligned each of them against the olive baboon amylase locus with NUCmer (v 3.1; default settings), visualising the results with mummerplot and extracting the corresponding coordinates. Between these species, uninterrupted collinearity was observed across the junctions that define the “first segment” and “fourth segment” as defined above. The “second segment” and “third segment” were then mapped separately against the ancestral two-copy configuration, allowing us to delineate the exact breakpoints introduced by the first and second non-allelic homologous recombination events. The sequence that is contiguous in olive baboons but split in the two-copy genomes corresponds to the novel duplications. These alignments confirmed that no additional rearrangements in the olive baboon amylase locus and that the derived *AMYp1* and *AMYp2* blocks are absent from the ancestral Catarrhini two-copy configuration, validating the BISER-based segmentation. We additionally aligned the olive baboon locus to that of its sister species Guinea baboon (*Papio papio*). Although the Guinea baboon assembly contains six tandem amylase copies, in addition to the ancestral *AMY2B* and *AMY1’* all of these extra copies are identical in nucleotide sequence, strongly suggesting an assembly artifact caused by assembly over-expansion rather than true structural variation in the locus (see **Methods**). Because Guinea baboons retain only the ancestral two-copy configuration, both *AMYp1* and *AMYp2* appear to be olive baboon-specific. Consequently, the two NAHR events that generated these duplications must have occurred after the complete split from Guinea baboons, within the ∼1.85 MYA window nferred for Papionini divergence

To investigate the macaque-specific duplication, we applied the same segmentation and alignment workflow to the rhesus macaque locus Self-alignments (dotplots) and pairwise NUCmer comparisons partitioned the region into three paralogue segments, each corresponding to a single duplicon: segment 1 harbouring *AMY2B* segment 2 carrying *AMYm* (the novel copy), and segment 3 harbouring *AMY1′* Pairwise comparisons showed that *AMYm* shares extensive 5′ homology with *AMY1′* and 3′ homology with *AMY2B* with crossover points falling within the intergenic sequence upstream the novel gene The preserved orientation and high identity of the flanks, together with these shared blocks, implicate NAHR as the duplication mechanism (**Figure S3**). n parallel, BISER (v. 1.4) identified 26 unique duplicons within the macaque locus (**Table S13**), which we treat as the smallest repeat units; by contrast, the three segments defined above represent the largest repeated modules spanning each paralogue block across the locus.

We then aligned orthologous contiguous amylase locus from multiple *Macaca* species representing the three major clades (fascicularis, sinica, silenus). *AMYm* is present in the *fascicularis* and *sinica* groups but absent from the *silenus* group, placing the duplication after the silenus split and before diversification of the fascicularis and sinica clade, i.e. approximately ∼4.5-5 MYA. All *AMYm*-harbouring loci were recovered on single, gap-free contigs with identical breakpoints across species and no additional rearrangements, supporting a single NAHR event that generated the three-segment architecture in macaques (**Figure S4**).

### Targeted Local Reassembly of the Guinea Baboon Amylase Locus

The original Guinea baboon (*Papio papio*) genome assembly (GCA_028858775.2) contained six identical tandem amylase copies in addition to *AMY2B* and *AMY1′* To evaluate whether this expansion reflects true structural variation or an assembly artifact, we adopted a targeted local reassembly approach. We extracted an extended bait region spanning from 150 kb upstream of the *RNPC3* 5′ flank to 150 kb downstream of the *AMY1C* 3′ flank, providing sufficient unique flanking context for unambiguous read recruitment. The full PacBio Revio HiFi read dataset (∼97 Gb) was mapped to this bait region using minimap2 (v. 2.29)^63^ retaining only primary alignments with mapping quality ≥20 and minimum aligned length ≥7 kb. Coverage was assessed in 20 kb windows across the extracted locus. A sharp drop in read depth was observed precisely over the region corresponding to the six tandem copies, inconsistent with a genuine six-copy expansion and instead indicative of assembly over-expansion.

To confirm this interpretation, the filtered reads were extracted and reassembled locally using both Flye (v. 2.9.6)^64^ and hifiasm (v. 0.25.0)^65^ Across mapping quality thresholds from MQ 10 to MQ 30 and minimum aligned lengths from 6 kb to 10 kb, hifiasm consistently produced a single continuous contig spanning the entire locus, containing only two amylase genes (*AMY2B* and *AMY1′*) corresponding to the ancestral Catarrhini configuration. Flye occasionally fragmented the locus at tandem repeat boundaries but ever supported more than two gene copies. As an additional control, the analysis was repeated using the gelada (*Theropithecus gelada*) amylase locus as an independent bait reference; uniform coverage and a two-copy local assembly were again recovered. Based o the locally reassembled locus, transposable element annotations for the Guinea baboon amylase region were recalculated using RepeatMasker with the same parameters applied to all other species, and al downstream analyses were updated accordingly to reflect the TE composition of the reassembled locus.

### Transcriptomic Data Generation and Processing and Differential Expression Analysis

Biopsies (parotid, submandibular, sublingual, pancreas and liver) from five olive baboons (*Papio anubis*) and six rhesus macaques (*Macaca mulatta*) were flash-frozen in liquid nitrogen immediately after collection and stored at -80°C. These samples were collected at Texas Biomedical Institute by veterinarians from 7-18 year old animals right after planned euthanization for health reasons Samples are kept at -20°C. Library preparation and standard Illumina HiSeq RNA sequencing (2x150bp) experiments were carried out following standard operating procedures by GENEWIZ/Azenta. Publicly available adult human salivary gland dataset^49^ was downloaded as FASTQ files and processed similarly to the olive baboon and rhesus macaque RNAseq samples from the quality-control step onwards. The pancreas and liver expression datasets were obtained from GTEx v8 and normalized as detailed below.

Quality assessment of raw reads was performed with FastQC (v. 0.12.1)^66^ and summarised with MultiQC (v. 1.25)^67^ Adapter sequences, low quality bases at the 3’ and 5’ end of the read, and reads shorter than 36 bp were removed using Cutadapt (v. 3.5)^68^ To investigate tissue-specific expression, trimmed reads were aligned to the olive baboon (*Papio anubis* Annotation Release 104) and rhesus macaque (*Macaca mulatta* Annotation Release 103) transcriptomes, and transcript abundances were quantified with Kallisto (v. 0.51.1)^50^ Transcript-level transcripts per million (TPMs) were summarized to the gene level. The gene-level quantifications were further normalized using the "lengthScaledTPM" method, for consistency in downstream analyses. These data were then filtered to include only high-confidence gene expression estimates (minimum of ten reads across all samples). The normalized gene expression profiles were used for differential expression analysis and visualization. For differential expression analysis, we utilized DESeq2 (v. 1.46.0)^69^ to compare gene expression profiles between tissues within species. We applied a Wald test to detect significant expression differences between groups. Genes with an adjusted p-value (FDR < 0.05) were considered significantly differentially expressed.

Human salivary gland reads^49^ were quantified with Kallisto against the NCBI *Homo sapiens* Annotation Release 110 transcriptome. Pancreas and liver read-count tables were obtained from GTEx v8 Before combining these data with the Kallisto-derived human salivary gland quantifications, we removed Ensembl version suffixes from every gene ID, retained only genes present in all samples and rounded Kallisto’s fractional estimates to integers so that every entry represented a aw count compatible with DESeq2. A DESeq2 object was then created using tissue as the design factor, and size factors were estimated with the default median-of-ratios procedure to generate library-normalised counts. Variance-stabilising transformation was applied for PCA-based quality control (GTEx pancreas GTEx liver or each salivary gland tissue) to confirm that batch effects did not dominate the variance structure. The resulting size-factor-normalised count matrix was used for all subsequent differential-expression analyses.

In parallel, we developed a polymorphism-aware pipeline to achieve higher resolution quantification of gene expression across tissues. This pipeline leverages nucleotide polymorphisms within RNA-seq reads to partition expression into transcript-specific contributions. We first aligned all human, rhesus macaque, and olive baboon transcripts, accounting for alternative splicing isoforms, to identify coding-sequence polymorphisms. For each species, we generated a single consensus FASTA sequence that captured the shared polymorphisms across the different transcripts within species. We then extracted RNA-seq reads previously mapped to each genome using HISAT2 (v. 2.2.1)^70^ retaining only those overlapping the amylase locus, and realigned them to the corresponding species-specific consensus FASTA with BWA (v. 0.7.19-r1273)^71^ NCBI annotations were curated for known polymorphic regions in the olive baboon and rhesus macaque transcripts, and the data were integrated with species-specific annotations to refine the analysis.

Using Old World Monkeys transcripts as outgroups, we assigned the detected polymorphisms as ancestral, derived, or lineage-specific. Using custom Python scripts we extracted and quantified reads at these polymorphic sites, enabling paralog-specific expression estimates (scripts have been deposited to Zenodo:https://doi.org/10.5281/zenodo.16809248). By focusing on transcript- and paralog-level variation, our approach aimed to reveal tissue-specific expression patterns that would otherwise remain obscured in conventional bulk RNA-seq data. To ensure robust detection of gene-level differences, we specifically utilized polymorphisms that distinguish one paralogous gene from another. In the rhesus macaque genome, for instance, three paralogous genes, *AMY2B AMYm* and *AMY1’* are annotated in that order. For each paralogous gene, we selected four polymorphisms: two located toward the proximal (5’) end and two toward the distal (3’) end of the coding region. At positions 93 and 111, *AMY2B* carries the polymorphisms G, T, *AMYm* carries A, G, and *AMY1’* carries A, T (**Figure S14**). At positions 737 and 753, *AMY2B* carries A, T, *AMYm* carries T, T, and *AMY1’* carries T, C respectively. The polymorphism-aware pipeline allowed us to unambiguously assign reads to individual paralogs and confirmed the accuracy of Kallisto’s expression estimates t was used only as an internal validation step and did not contribute to the differential expression analyses or any other downstream analyses reported here.

Finally, we queried all annotated long-non-coding RNAs located within, or immediately flanking, the amylase locus in the rhesus macaque and olive baboon genome builds. The Kallisto-derived TPMs for every such lncRNA were below ten reads in every tissue, indicating negligible expression; they therefore cannot account for the tissue-specific patterns described in Results.

### ddPCR-Based Copy Number Estimation in Individuals Included in the Transcriptomics Analysis

To determine whether within-species variation in amylase copy number could confound paralog-specific expression comparisons, we quantified genomic amylase copy number by droplet digital PCR (ddPCR) in the same five olive baboons (*Papio anubis*) and six rhesus macaques (*Macaca mulatta*) for which RNA-seq data were generated from parotid, submandibular, sublingual, pancreas, and iver biopsies (see RNA-seq sample description above). Genomic DNA was isolated from each individual and analyzed using ddPCR, targeting a sequence conserved across amylase gene homologs in Old World monkeys. Copy number was calculated relative to a single-copy reference locus used in our previous work^17^ (**Table S14**). The assay used the following oligonucleotides: forward primer (5′-3′) GAGCACTTGTCTTTGTGGATAA; everse primer (5′-3′) TCCCAGAAGGTAAGAATAGAGG; and hydrolysis probe (5′-3′) CCATGACAATCAACGAGGACATGGG. Because the target region is shared among paralogs, this approach provides an aggregate estimate of total amylase copy number and does not resolve paralog-specific contributions to copy number variation.

### Positive Selection Analyses

Coding sequences (CDS) from the two Old World monkey species analysed here (olive baboons and rhesus macaques) together with orthologous CDS from all extant Great Ape species (human, chimpanzee, bonobo, Sumatran and Bornean orangutan) were aligned at the amino-acid level with MAFFT (v. 7.515)^72^ back-translated to codons with PAL2NAL (v. 14)^73^ and manually trimmed to preserve reading frame. We assessed the sequences for internal stop codons and removed any truncated CDS. A maximum-likelihood gene tree was inferred with IQ-TREE (v 2.4.0)^74^ using the MG+F3X4+R2 model and was input to all HyPhy analyses. All selection analysis results are provided in Supplementary **Tables S15-S18**

### Branch-level tests (aBSREL)

To identify entire lineages that experienced bursts of adaptive change we ran the adaptive Branch-Site Random-Effects Likelihood (aBSREL) model in HyPhy (v. 2.5.48)^75^ with default settings. aBSREL compares for every branch, a null model in which all sites evolve neutrally or under purifying selection (ω ≤1) to an alternative model that allows a proportion of sites on that branch to have ω>1 Likelihood-ratio tests are corrected for multiple comparisons with the built-in Holm procedure. Foreground branches were not pre-specified; instead HyPhy evaluates each branch in turn. The aBSREL p-values were corrected for multiple testing with HyPhy’s built-in Holm-Bonferroni procedure (**Table S15**).

### Site-level episodic diversifying tests (MEME)

Codon-specific episodic diversifying selection was evaluated with the Mixed-Effects Model of Evolution (MEME) in HyPhy^76^ MEME allows the selective regime (ω) at a site to vary among branches, thus detecting positive selection that operates only on a subset of lineages. We used the default significance cut-off of p≤0.10 after applying the False Discovery Rate (FDR) correction using the Benjamini-Hochberg procedure (**Table S16**).

### Site-level pervasive tests (FUBAR)

Pervasive, site-wide selection was assessed with the Fast, Unconstrained Bayesian AppRoximation (FUBAR) implemented in HyPhy^77^ which estimates posterior probabilities for ω>1 under a Bayesian framework that assumes a constant selection regime across the tree. Analyses were run for 5 million MCMC iterations (burn-in=1 million); codons with posterior probability larger than 0.90 were considered positively selected (**Table S17**).

### Relaxation or intensification of selection (RELAX)

To test variation in selective pressures across specific lineages, we applied RELAX^78^ The baboon *AMYp2* branch was assigned as foreground and all other branches as background. RELAX fits two models differing by a scaling parameter K that inflates (K>1) or deflates (K<1) the background ω distribution; significance is assessed by a likelihood-ratio test (**Table S18**).

### Structural Modeling and Functional Domain Prediction

The coding sequence for *AMYp2* was modelled *in silico* using AlphaFold2 (v. 2.3.1)^79^ via the ColabFold implementation with the “monomer_ptm” preset and default recycling.The top-ranked model by pLDDT was retained; predicted-aligned-error (PAE) matrices were inspected to verify global fold confidence. Annotated PDB files were generated in Chimera (v. 1.19) and used for all subsequent structure-based alignments.

Active, catalytic and calcium-binding sites were identified using the NCBI Conserved Domain Database (CDD, accessed March 2025). Calcium and chloride binding sites were further cross-referenced with the identified binding sites by Ramasubbu et al.^80,81^ Glycosylation candidates were predicted with NetNGlyc 1.0^82^ and cross-referenced to the proposed glycosylation sites reported by Kamitaki et al.^45^

### Regulatory Motif Analysis

Promoter sequences for the amylase paralogs in humans and rhesus macaques were defined as the 170-bp window spanning 100 bp upstream to 70 bp downstream of the experimentally supported TSS recorded in Eukaryotic Promoter Database (EPD)^83^ When an EPD entry was unavailable, we took the RefSeq transcription-start site (TSS) from the corresponding annotation (NCBI *Papio anubis* Release 104, *Macaca mulatta* Release 103, and *Homo sapiens* Release 110) and defined the promoter as the region 100 bp upstream to 70 bp downstream of that TSS. We retrieved these windows for every amylase paralog in humans, rhesus macaques, and olive baboons. In parallel, we analyzed the 50 most highly expressed salivary gland genes in each species (ranked by TPM) and annotated their promoter sequences. Each promoter was scanned with MEME (MEME-suite v. 5.5.7)^84^ for *de novo* motif discovery to identify novel, enriched sequence motifs.

The identified motifs were then annotated with Tomtom (MEME-suite v. 5.5.8)^85^ against the JASPAR 2024 CORE non-redundant vertebrate library^86^ retaining hits with P<10^-4^ to identify potential transcription factors (TFs). To assess specificity, we conducted parallel analyses using promoter sequences from 50 randomly selected genes per species, which did not show significant motif enrichment for the motifs identified with the salivary gland dataset, supporting the specificity of our results The resulting TF list was cross-referenced with salivary- and pancreas-specific transcription factors with enriched expression from the FANTOM5 database (P<0.05 and log_10_(relative expression over median)>1.3)^87^ and the salivary gland TF catalogue from Michael et al.^51^ TFs present in either salivary gland set and absent from the pancreatic set were labelled salivary-gland-biased, while the converse defined pancreatic-biased TFs, with the remainder classed as core The amylase-paralog promoter windows were then scanned with FIMO^88^ against the JASPAR 2024 CORE library to identify TFBS within each promoter; only hits with P<10^-4^ were retained. These TFBS were subsequently grouped into core pancreatic-biased, and salivary-gland-biased categories based on the above TF assignments.

## ADDITIONAL RESOURCES

Analysis scripts, pipelines and input data: https://doi.org/10.5281/zenodo.16809248 and https://doi.org/10.5281/zenodo.18689074

## Notes

### Competing Interest Statement

The authors have declared no competing interest.

### Summary of Updates

Methods and Limitations have been updated.

### Funding

- NHI: R35-GM156519
- National Science Foundation: 2049947, 2123284
- U.S. National Science Foundation: PRFB. 2508185

