## Supplemental Data 1 for "Convergent evolution through independent rearrangements in the primate amylase locus"

### GLOSSARY

**Segmental Duplications:** Segmental duplications (SDs) (also called low-copy repeats, LCRs) are stretches of DNA present two or more times per haploid genome, with each copy exhibiting a high degree of sequence identity (commonly >95%) and typically spanning at least ~1 kilobase in length. These duplicated segments may occur in tandem (*i.e.* adjacent to one another) or may be dispersed to different locations on the same chromosome or on different chromosomes.

**Recurrent Duplications:** Recurrent duplications refer to duplication events that arise multiple times, independently, at the same genomic locus across different individuals or lineages. These events are typically mediated by underlying genomic features, such as the presence of flanking segmental duplications or low-copy repeats or transposons or mutational hotspots, which predispose these loci to recurrent rearrangement through mechanisms like non-allelic homologous recombination (NAHR).

**Independent Duplications:** Independent duplications denote duplication events that have arisen separately, (not from a shared ancestral event) as separate mutational events, either at the same or different loci. In this context, "independent" emphasizes that each duplication occurred as a unique mutational event, not inherited from a common ancestor.

**Recurrent Independent Duplication:** Recurrent independent duplications are structural rearrangements in which the same genomic locus is duplicated multiple times, but each duplication arises as a distinct, independent mutational event in different lineages or individuals, rather than being inherited from a common ancestor.

- Recurrent: The duplication happens repeatedly at a specific locus, often due to the region's inherent genomic instability or predisposition to rearrangement.
- Independent: Each observed duplication event is phylogenetically separate i.e. arising de novo in different evolutionary lineages or individuals, rather than being a single ancestral duplication that is inherited.

**Lineage-specific events:** Lineage-specific events are genetic or evolutionary changes, such as deletions, duplications, regulatory shifts etc, that occur uniquely within a particular evolutionary lineage after it has diverged from its closest relatives. In phylogenetic terms, these are changes that take place along the branch (edge) connecting two nodes in a species tree. Such events distinguish that lineage and the species it contains from all others due to novel mutational or biological processes that arose within that lineage alone.

**Convergent Evolution:** The independent emergence of similar traits or features in related lineages that had different ancestral starting points. In convergence, similar phenotypes arise via different genetic changes or developmental routes, but are shaped by similar ecological or selective pressures.

**AMY:** Alpha-amylase gene family: A family of genes encoding alpha-amylase enzymes, which hydrolyze starch into sugars. Members of this family often arise via gene duplication and can differ in tissue specificity, expression level and function.

***AMY1/AMY1A|B|C*:** Great ape salivary gland amylase genes: Salivary gland-specific amylase genes in great apes, derived from the ancestral *AMY1'* gene, primarily involved in starch digestion in the oral cavity.

***AMY2A*:** Great ape pancreatic amylase gene: A pancreatic-specific amylase paralog in great apes, evolved from the same the ancestral *AMY1'* gene that gave rise to *AMY1*, and specialized for starch digestion in the small intestine.

***AMY2B*:** The presumed ancestral amylase gene in primates, with conserved expression in the pancreas across multiple lineages.

***AMY1'*:** Ancestral Catarrhini amylase duplicate: The first duplication of *AMY2B* in the Catarrhini ancestor, predating the split between Old World monkeys and apes; progenitor of human *AMY1* and *AMY2A*.

***AMYm*:** Macaque-specific amylase duplicate: A lineage-specific duplication of *AMY2B* in macaques (fascicularis and sinica groups), arising after divergence from the silenus group.

***AMYp1*:** Baboon-specific first amylase duplication: The first baboon-lineage duplication generated via non-allelic homologous recombination of the *AMY2B* and *AMY1'* paralogs, shared with the Guinea baboon.

***AMYp2***: Olive baboon-specific second amylase duplication: A second, more recent duplication in the olive baboon lineage, derived from *AMY2B* and *AMYp1* via non-allelic homologous recombination.

**ABBREVIATIONS**

**OWM:** Old World monkey

**NWM:** New World monkey

**NAHR:** Non-allelic homologous recombination

**TFBS:** Transcription factor binding sites

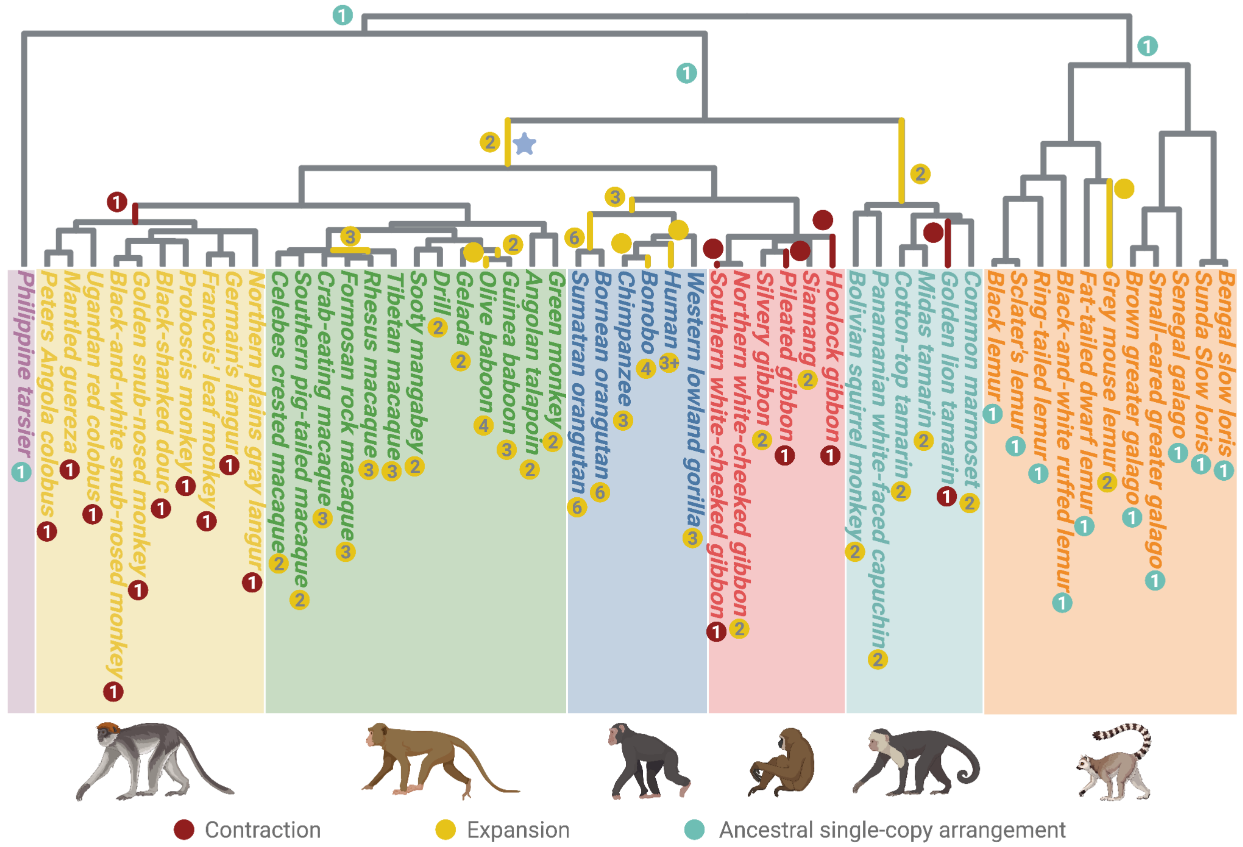


**Figure S1.** Evolutionary dynamics of the amylase locus across primates. This figure reproduces the phylogeny shown in Figure 1A but replaces Latin names with common species names to improve readability. In addition, inferred amylase copy numbers are annotated along non-terminal branches based on parsimony reconstruction. The Catarrhini lineage is marked with a purple star. Contractions and expansions of the amylase locus reconstructed from 69 high-quality genomes representing 53 primate species. Clades are color-coded as follows: purple, tarsiers (outgroup); yellow, colobine leaf-eating monkeys; green, cercopithecine Old World monkeys; dark blue, great apes; red, lesser apes (gibbons); cyan, New World monkeys; orange, lemurs. Red circles indicate independent contraction events; yellow circles indicate independent expansions; cyan circles denote lineages retaining the inferred ancestral single-copy configuration. Numbers within circles represent total AMY gene copies per haploid genome. Numbers along internal branches denote inferred copy-number changes occurring on non-terminal branches.

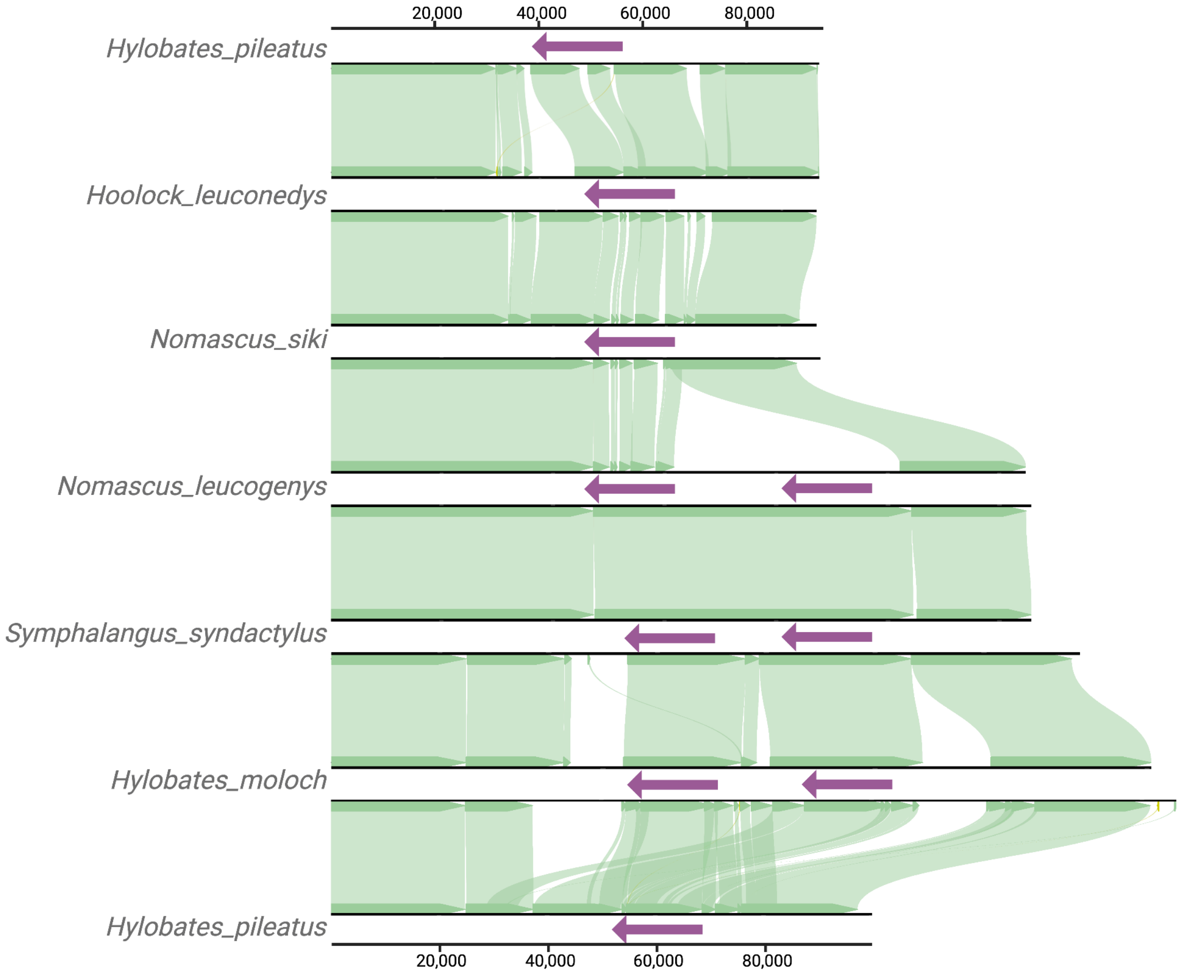


**Figure S2.** Copy-number and structural diversity of the amylase locus in gibbons (lesser apes). Miropeat style plots across representatives of *Hoolock leucodenys*, *Symphalangus syndactylus*, *Hylobates pileatus, Hylobates moloch* and *Nomascus leucogenys* and *Nomascus siki* illustrate lineage-specific differences consistent with either independent losses or incomplete lineage sorting (see main text). Axes denote locus coordinates.

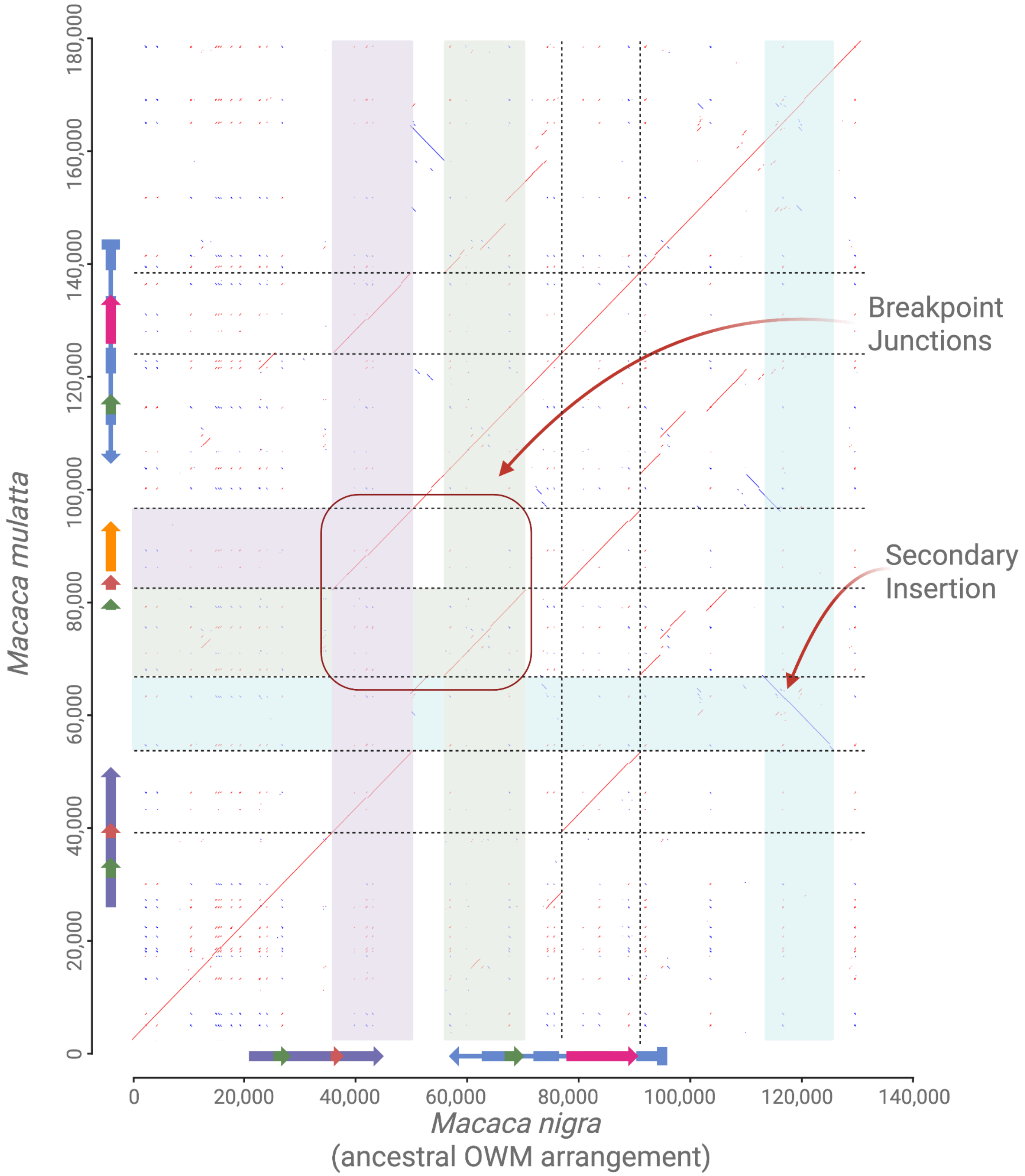


**Figure S3.** Synteny of the macaque amylase locus across *Macaca* and mapping of the *AMYm* duplication. Comparative dotplots use *Macaca nigra* as the two-copy ancestral Old World monkey configuration and highlight the novel *AMYm* block in the fascicularis and sinica clades (represented by *M. mulatta*), with breakpoints marked at the NAHR junction and a small secondary insertion within the novel duplicon.

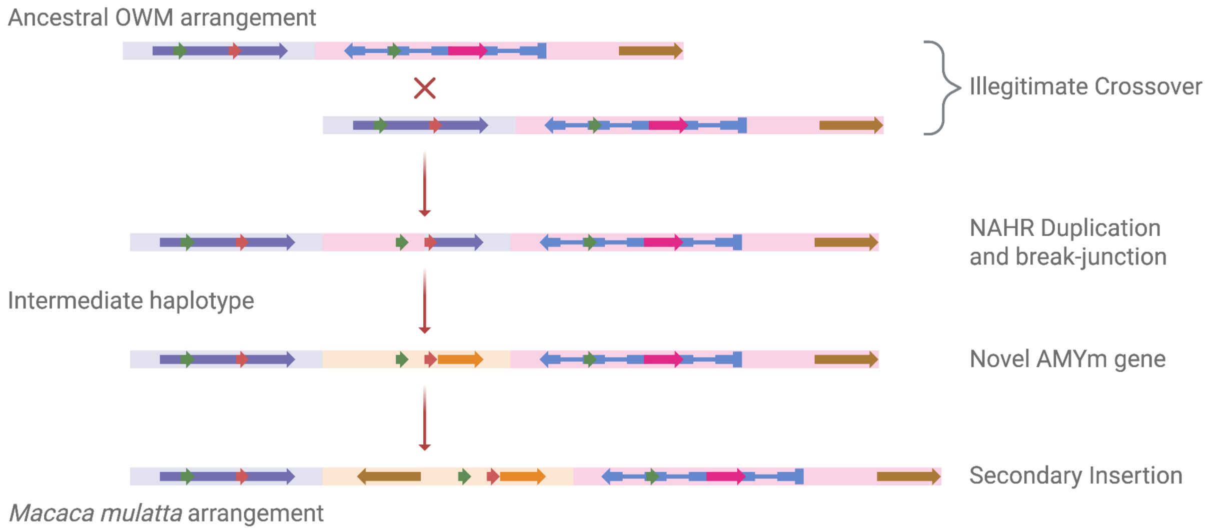


**Figure S4.** Mechanistic model for the macaque *AMYm* origin. Cartoon summarizing an initial illegitimate crossover (NAHR) between *AMY1′* (5′ homology) and *AMY2B* (3′ homology), generating the chimeric *AMYm* (intermediate haplotype). Following the NAHR duplication the locus experienced a small secondary insertion upstream the *AMYm*. Panels show the ancestral OWM configuration, *Macaca mulatta* arrangement, and the inferred intermediate arrangement prior to the secondary insertion.

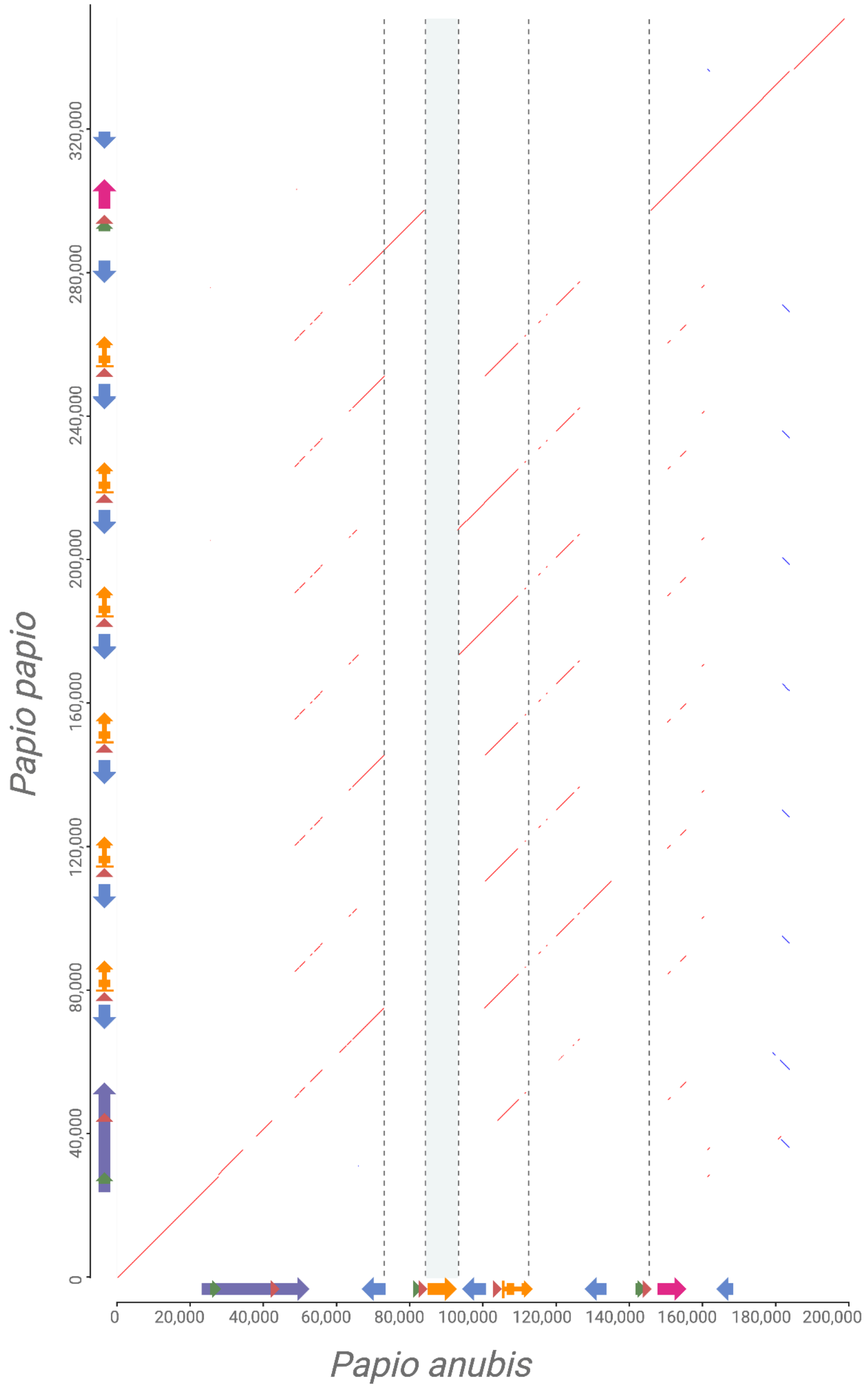


**Figure S5.** Dotplot of the amylase locus between *Papio anubis* (olive baboon; x-axis) and *Papio papio* (Guinea baboon; y-axis). *P. papio* lacks the *AMYp2* segment present in *P. anubis*, confirming *AMYp1* as the first and older duplication shared by both species. In *P. papio*, multiple tandem tracts align identically to *AMYp1* (6 copies), a pattern consistent with assembly over-expansion rather than a true biological increase (see Methods). Alignments were generated with LAST (using lastal) and filtered with last-split to retain only primary, best-scoring placements and the dotplot was rendered from these primary chains.

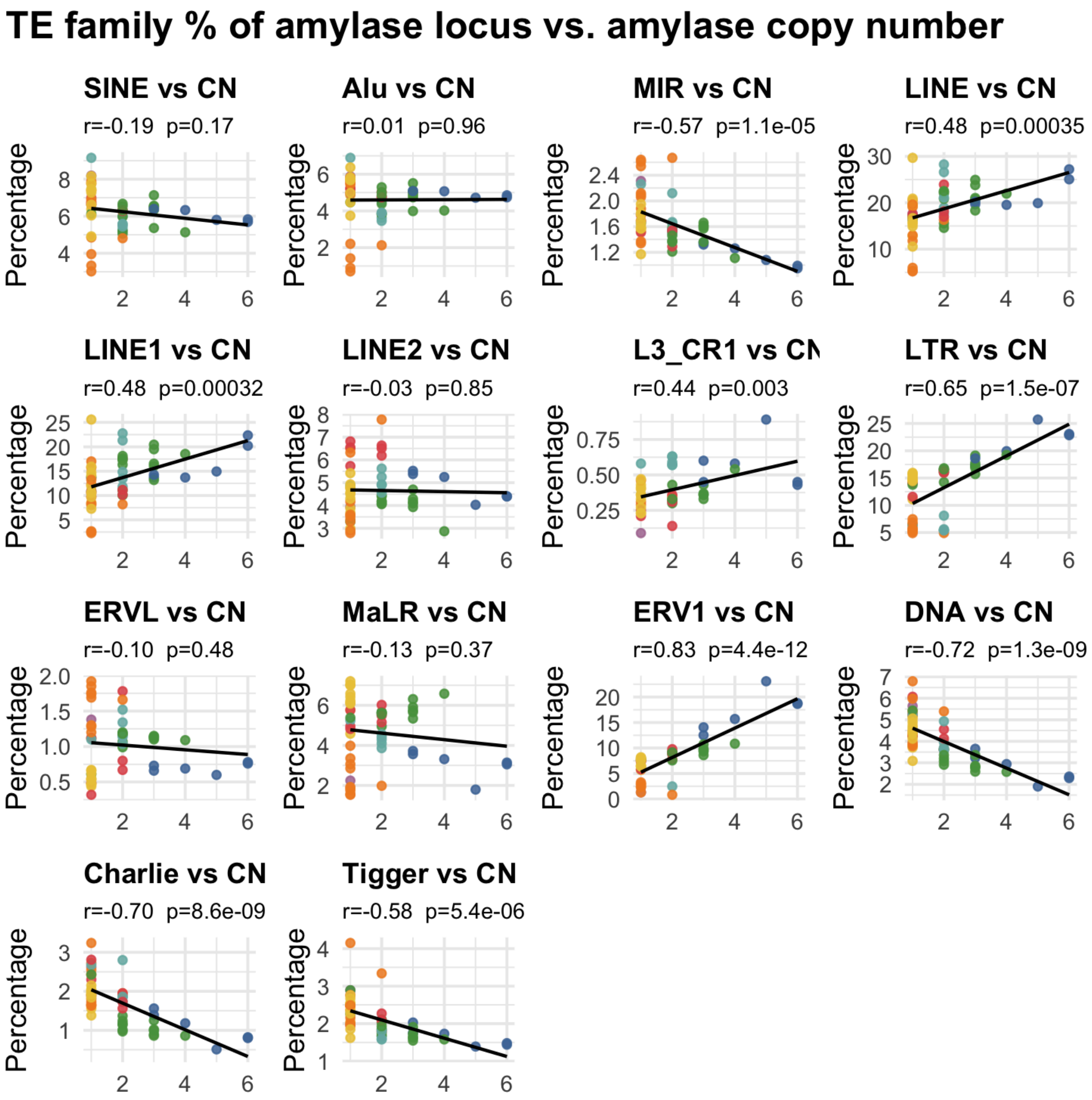


**Figure S6.** TE family abundance in the amylase locus versus AMY copy number across primates. Scatterplots show, for 53 species, the percentage of each TE family within the RNPC3-AMY1C amylase interval (y-axis) against the total AMY copy number per haploid genome (x-axis). Points are species (colored by clade as in Fig. 1); lines are least-squares fits. Panels report Pearson's r and two-sided P test. LTRs, LINEs (LINE1, L3/CR1) and ERV1 show positive correlations with AMY copy number, whereas MIRs and DNA transposons (Charlie, Tigger) are negatively correlated. TE content was called with RepeatMasker using a uniform primate library on locus-bounded sequences.

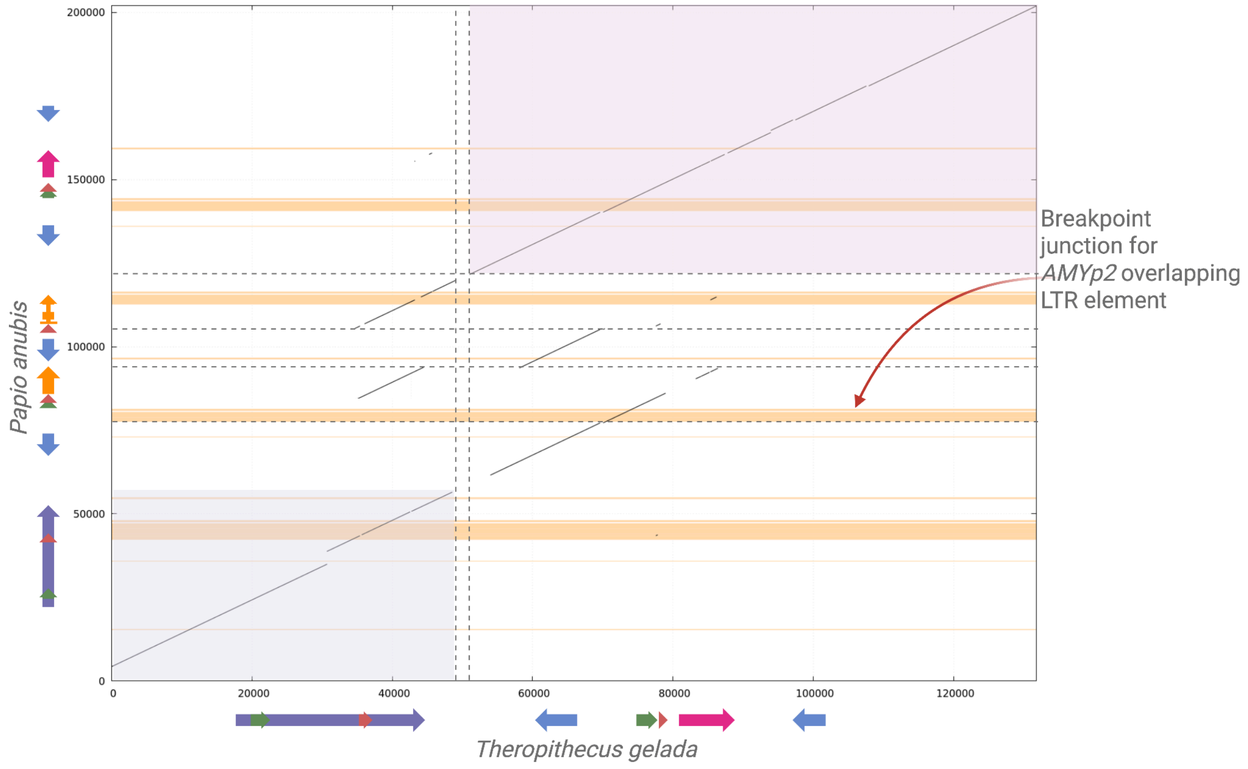


**Figure S7.** Dotplot of the olive baboon (*Papio anubis*) amylase locus (y-axis) versus the inferred ancestral Old World monkey two-copy locus from gelada (*Theropithecus gelada)* (x-axis), aligned with LAST (primary alignments only). Light purple background blocks mark the two ancestral OWM segments. Horizontal orange bands are LTR annotations in olive baboons. The dashed horizontal lines mark the inferred crossover positions. The labeled *AMYp2* breakpoint junction overlaps an LTR tract (LTR25-int).

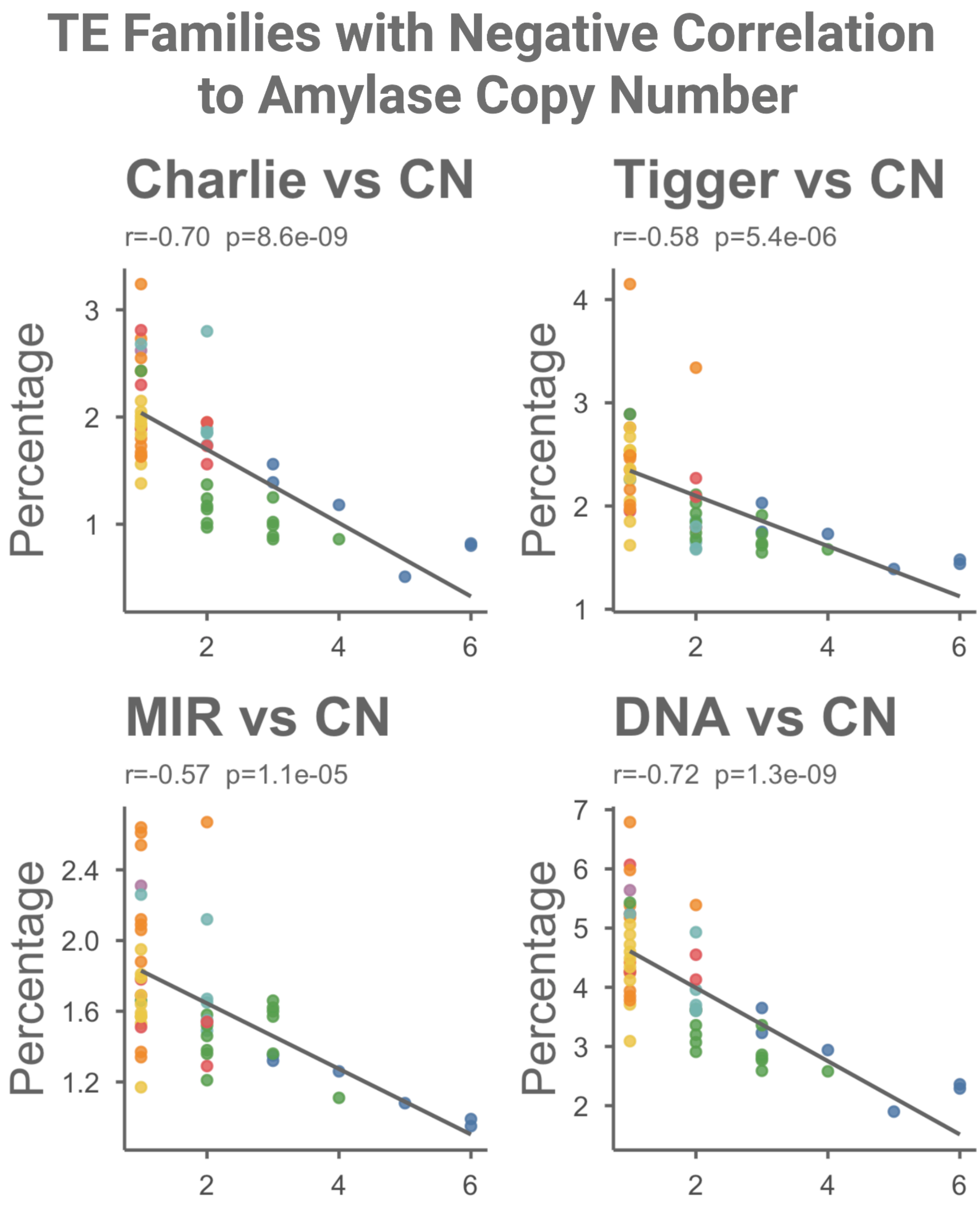


**Figure S8.** TE families negatively correlated with AMY copy number. Zoomed scatterplots for the four families with significant negative associations in Figure S6 (Charlie, Tigger, MIR, and total DNA transposons). Axes, point colors, fit lines, and statistics are as in Figure S6.

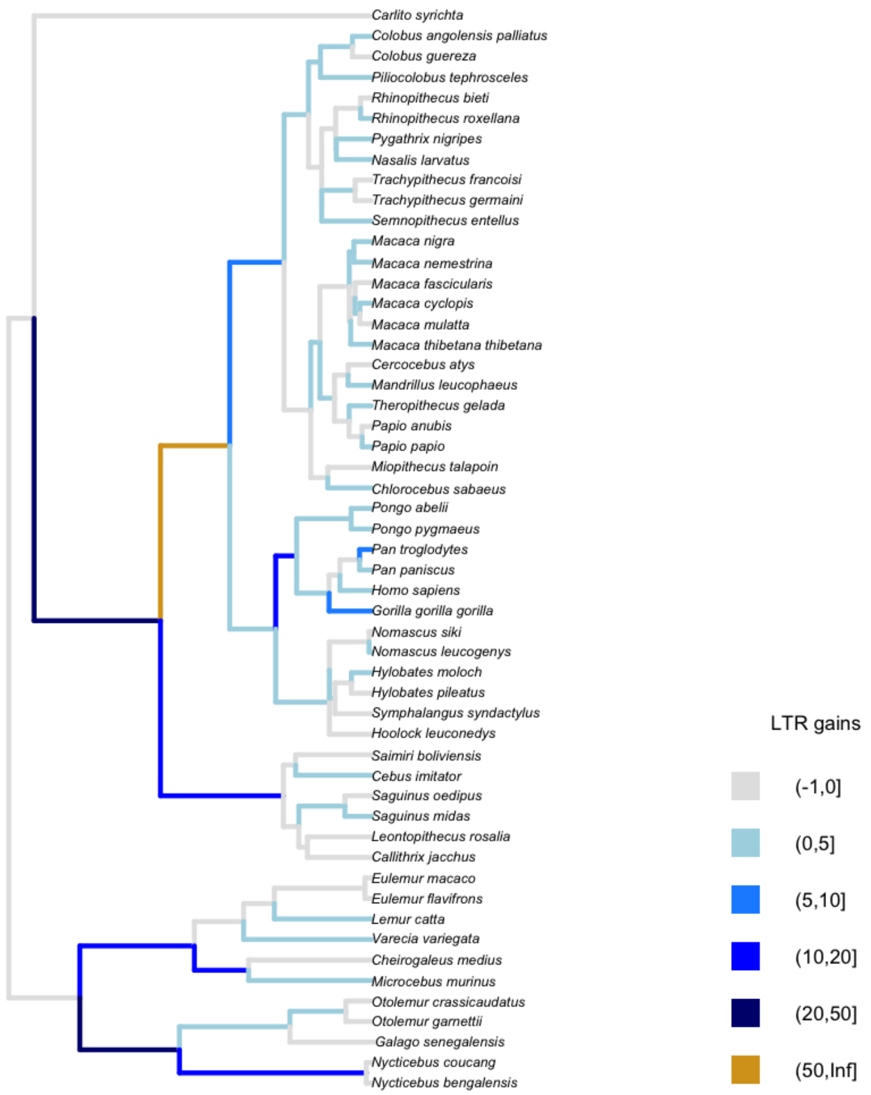


**Figure S9.** LTR gains at the amylase locus across primates. Branch-wise reconstruction of LTR insertion gains at the amylase locus across 53 primate species. Orthologous LTR insertions were identified using a reciprocal 1kb flanking-sequence BLAST approach and clustered into 214 orthogroups. A binary presence/absence matrix was analyzed on the pruned primate phylogeny. Ancestral states were reconstructed using maximum-likelihood discrete-character analysis under an equal-rates (ER) model and gains were inferred by comparing “parent-child” state transitions across all edges and orthogroups. Branch colors are proportional to the total number of inferred LTR insertion gains per branch, with darker shades indicating higher counts.

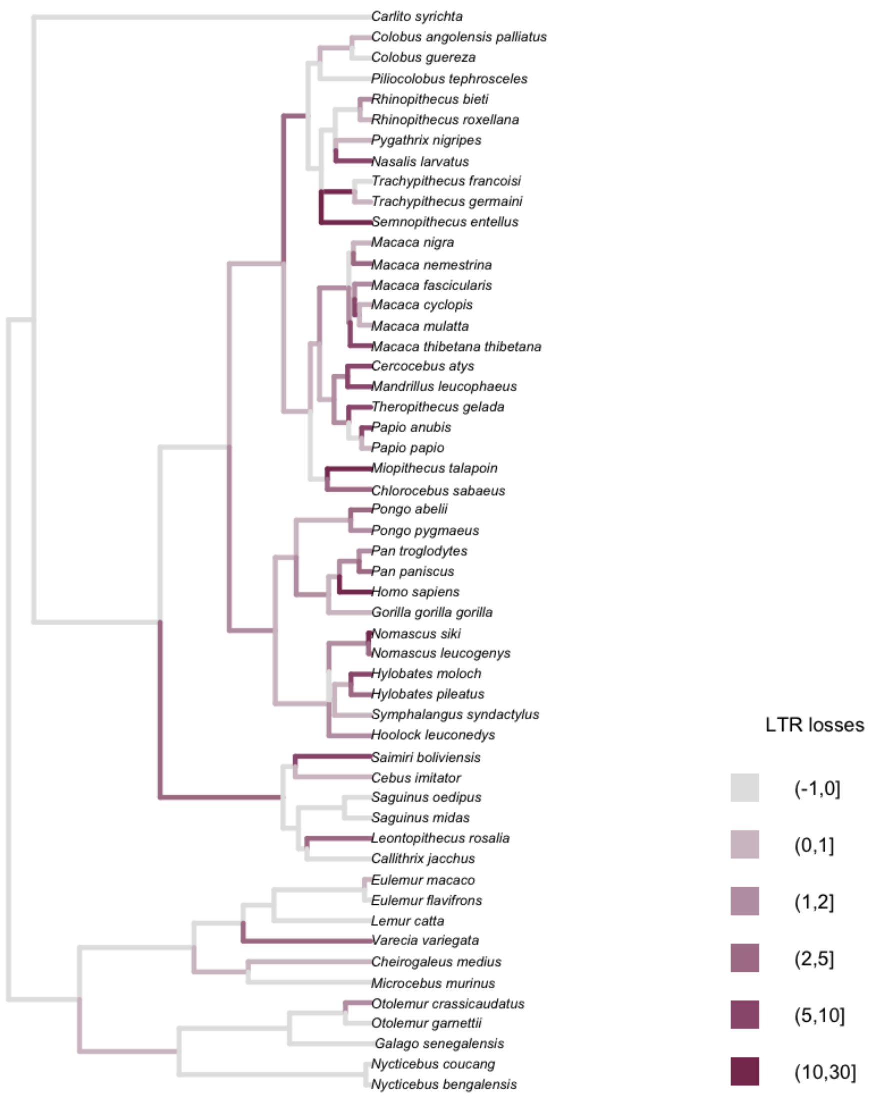


**Figure S10.** LTR losses at the amylase locus across primates. Branch-wise reconstruction of LTR insertion losses at the amylase locus across 53 primate species. Orthologous insertions were defined using reciprocal flanking-sequence comparisons and clustered into 214 orthogroups. Maximum-likelihood ancestral state reconstruction under an equal-rates (ER) model was used to infer state transitions along the phylogeny. Losses were quantified by comparing "parent-child" states across all branches and orthogroups. Branch colors represent the total number of inferred LTR losses per branch, with darker shading indicating greater numbers of loss events.

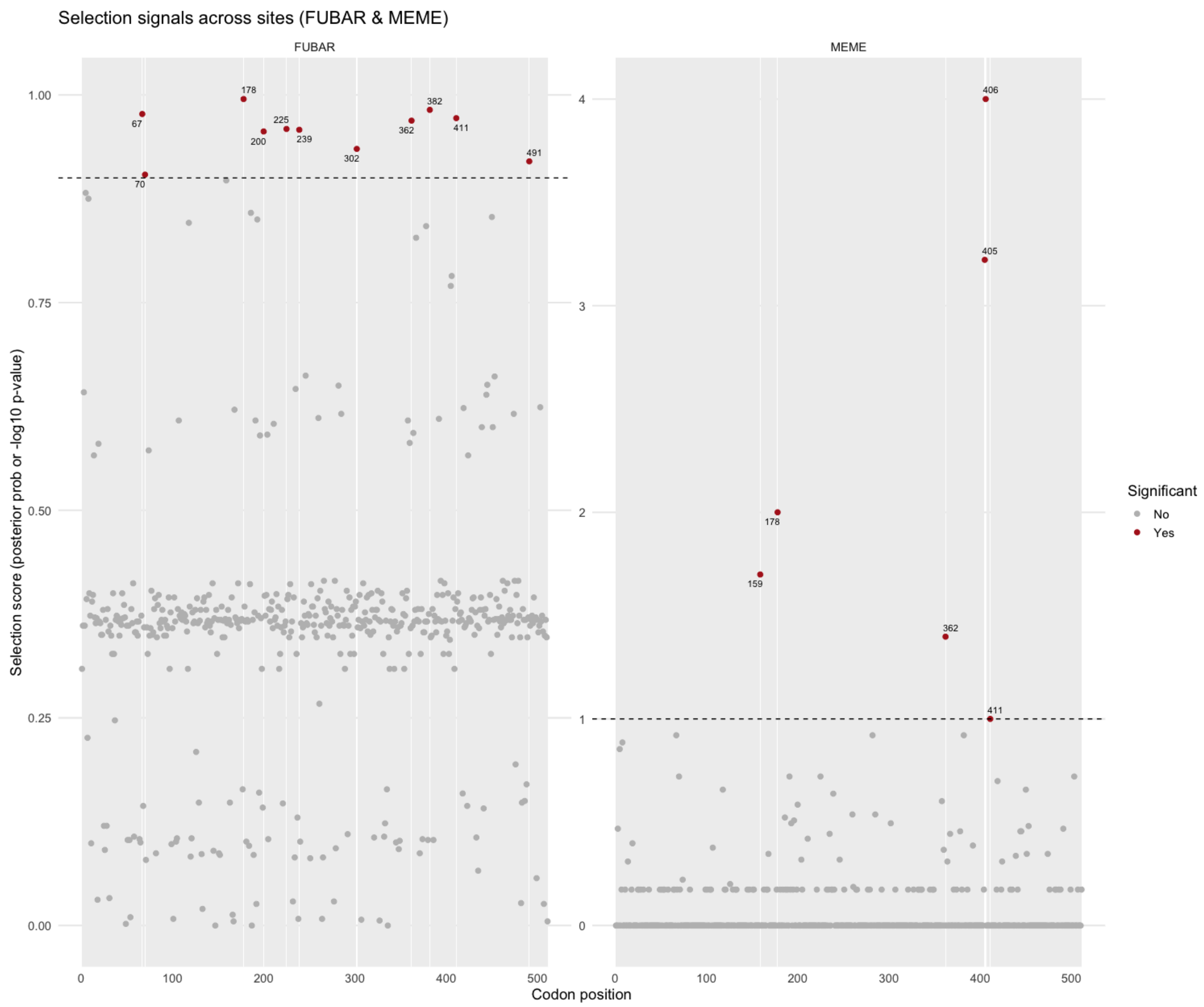


**Figure S11.** Site-level signals of selection across amylase coding sequences. Left: FUBAR posterior probability that ω>1 at each codon; the dashed line marks the 0.90 significance threshold. Right: MEME $-\log_{10}(P)$ for episodic selection; the dashed line marks P=0.10. Red dots denote sites called significant by each method (e.g. MEME: 159, 178, 362, 405, 406, 411; FUBAR: multiple sites ≥0.90). Analyses were run on the codon alignment of Old World monkey and great ape *AMY* paralogs.

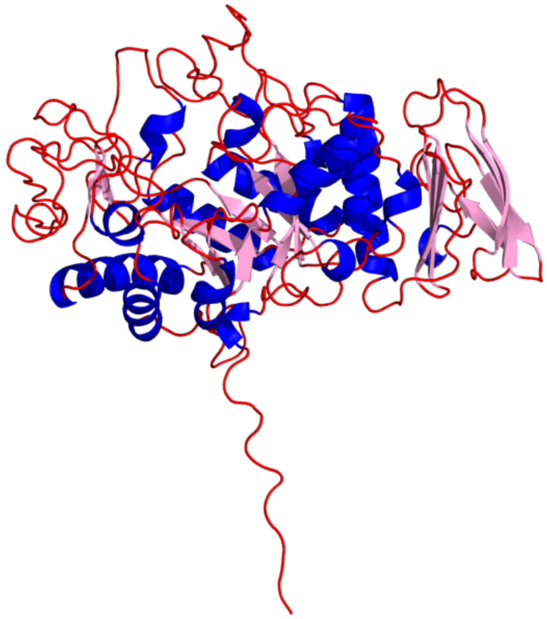

**Figure S12.** Predicted 3D structure of olive baboon *AMYp2*. AlphaFold2 model rendered as a cartoon with α-helices in blue, β-strands in light pink, and loops/coil in red.

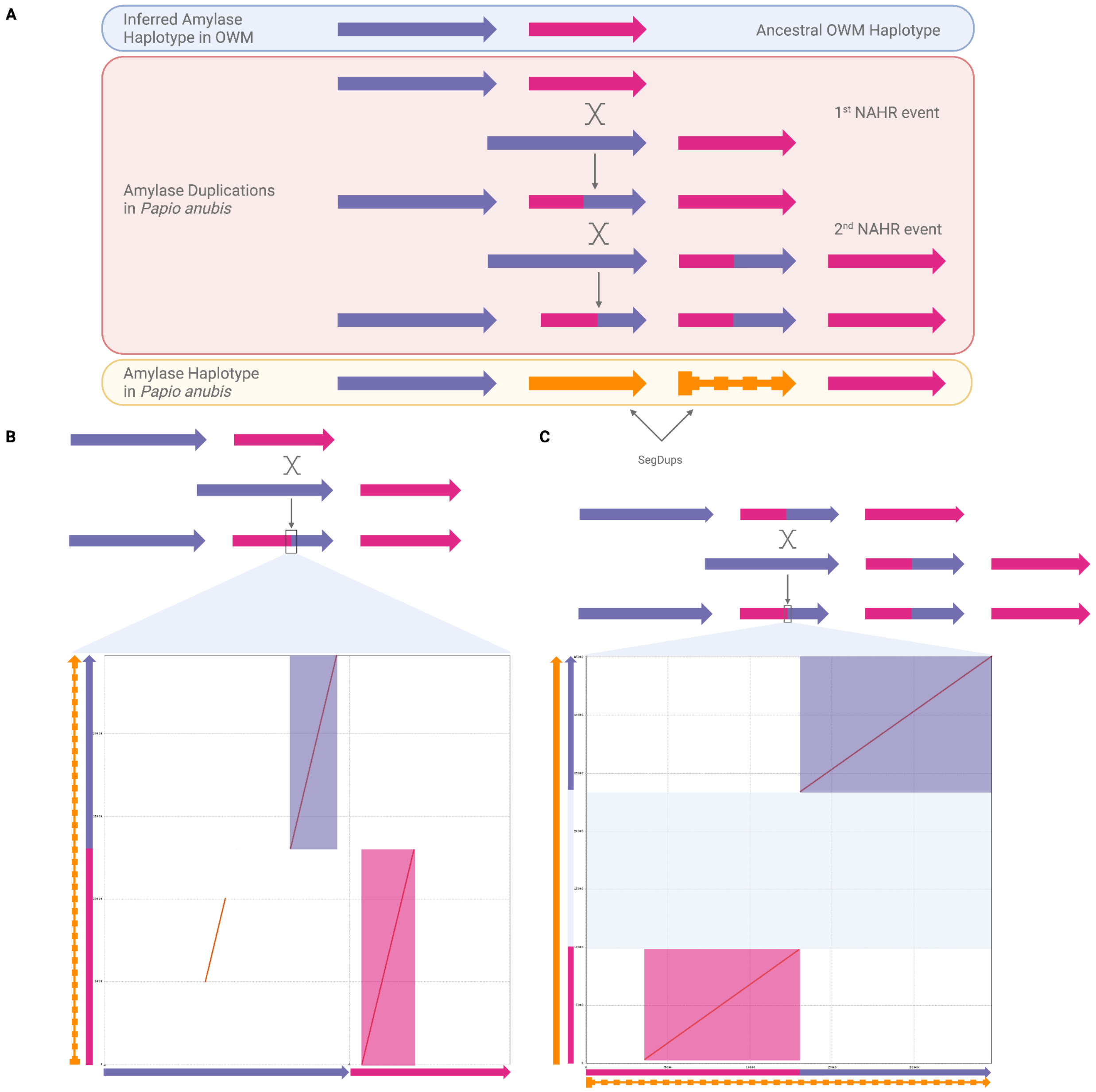


**Figure S13.** Stepwise NAHR duplications shaping the olive baboon amylase locus. The locus is partitioned into four duplicated segments, each drawn as a colored arrow to indicate block identity and orientation; colored arrows denote segments, not individual genes. Segment 1 (purple arrow) carries *AMY2B*, segment 4 (pink arrow) carries *AMY1'*, segment 3 (orange-dashed arrow) carries *AMYp1*, and segment 2 (orange arrow) harbors *AMYp2*. (A) Pairwise NUCmer alignments and dotplots show segment 3 is a mosaic of segment 1 and segment 4, consistent with a first NAHR event between segments 1 and 4 that created the *AMYp1*-containing block. (B) Segment 2 ( which hardbours *AMYp2*) is near-identical to segment 3, except for an ~10 kb DNA stretch absent from segment 3. Because this interval is flanked by long N stretches in the olive baboon assembly, we cannot determine whether it reflects a true deletion or a scaffolding artifact. These dotplots and cartoon schematic representation support a two-step NAHR model: NAHR between segments 1 and 4 produced segment 3 (harboring *AMYp1*), followed by NAHR between segment 1 and segment 3 produced segment 2 (harboring *AMYp2*), yielding the present-day order *AMY2B* *AMYp2*, *AMYp1*, *AMY1'* in olive baboon. Segments were defined from self-alignments/BISER and compared with NUCmer dotplots as described in Methods.

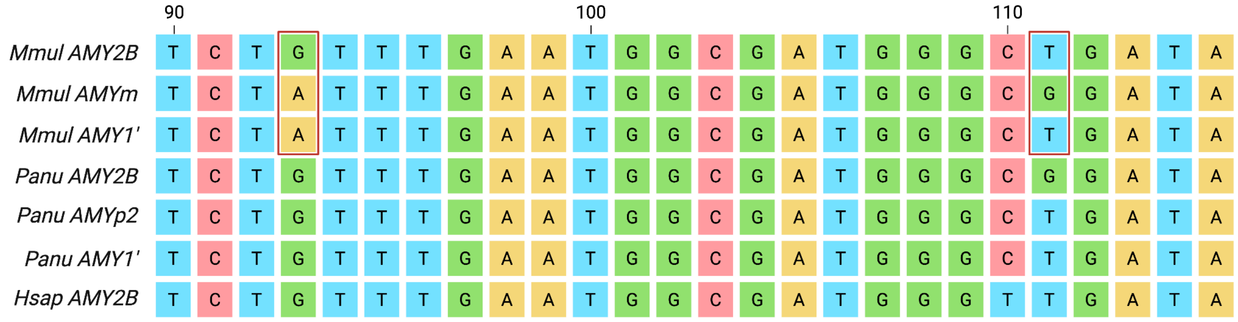


**Figure S14.** Example of the diagnostic polymorphisms used for paralog-specific expression in macaque and baboon. Alignment snapshot (codons ~90-10) across AMY paralogs shows fixed nucleotide differences used to assign RNA-seq reads to specific copies. Red outline boxes highlight parsimony informative sites in macaque (e.g. positions 93 and 111 distinguishing *AMY2B*, *AMYm*, and *AMY1'*). These markers underlie the SNP-aware quantification described in Methods.
